# Multiple Glycans and IFITM Proteins Mediate Human Coronavirus OC43 entry

**DOI:** 10.64898/2026.11.18.767684

**Authors:** Manivel Lodha, Masashi Iwamoto, Michael Tartell, Marie Canis, Noah A. Schuster, Daniel Poston, Yiska Weisblum, Stanley Perlman, Theodora Hatziioannou, Paul D. Bieniasz

**Author notes:** Authors contributed equally (Manivel Lodha, Masashi Iwamoto).

## Abstract

Human coronavirus (HCoV)-OC43 is the most prevalent seasonal coronavirus, causing colds, and occasional severe disease. HCoV-OC43 attachment is reportedly mediated by 9-O acetylated sialic acid (9-O-Ac-Sia); no proteinaceous receptor has been identified. Using genome-wide CRISPR-cas9 screens, we identify several host factors that mediate HCoV-OC43 entry. First, we identify NXPE3, an esterase, as required for HCoV-OC43 attachment and infection, both *in vitro* and *in vivo*, and demonstrate a previously unknown role for NXPE3 in 9-O-Ac-Sia synthesis. However, we further show that single residue substitutions in HCoV-OC43 spike, in the S1A domain at the 9-O-Ac-Sia binding site, or two substitutions in the S1B domain, confer 9-O-Ac-Sia independence and a switch to heparan sulfate-dependent entry. Additionally, we find that IFITM1/2/3 proteins act post-attachment to drive HCoV-OC43 virion-cell fusion and entry, irrespective of which attachment mechanism is used. Thus, flexible use of glycan attachment factors enables HCoV-OC43 to access fusion-driving IFITM proteins during entry.

## Main

Human coronavirus OC43 (HCoV-OC43) is an enveloped single-stranded positive-sense RNA virus of the Beta coronavirus genus (*Embecovirus* subgenus). HCoV-OC43 was isolated in 1967 from patients with respiratory disease [1] and is closely related to Bovine coronavirus (BCoV), sharing 96% sequence identity. Molecular clock analysis suggests that HCoV-OC43 zoonotic transmission, from cattle to humans, likely took place at around 1890, [2]. HCoV-OC43 is the most common cause of coronavirus upper respiratory tract infections and is associated with neurovirulence in humans and animal models [3]. With the emergence of SARS-CoV [4], MERS-CoV [5] and most recently, SARS-CoV-2 [6, 7] it is clear that coronaviruses have a severe impact on humans, highlighting the importance of understanding their biology.

Coronaviruses spike proteins mediate entry and are type-I transmembrane proteins that are processed into S1 and S2 subunits and form a trimer of S1/S2 dimers. S1 subunits form the outer ‘head’, with two globular regions termed the S1 N-terminal domain (S1 NTD/S1A) and S1 C-terminal domain (S1 CTD/S1B). The S2 domain forms a ‘stalk’ comprising the fusion machinery, including a membrane-anchoring transmembrane domain and cytosolic tail [8]. Coronaviruses’ spike proteins engage cell-surface glycans and/or protein receptors through their receptor-binding domain (RBD), usually present in the spike CTD/S1B [9, 10]. Receptor interaction, and spike cleavage by cell surface proteases (TMPRSS2) [11–15] or endosomal cathepsins [14, 16, 17] triggers conformational changes in spike, exposing a fusion peptide that inserts into cell membranes to initiate fusion. With the exception of HCoV-OC43, protein receptors have been identified for all human coronaviruses, namely ACE-2 (SARS-CoV [18], HCoV-NL-63 [19], SARS-CoV2 [20]), DPP4 (MERS-CoV) [21], APN (HCoV-229E) [22] and TMPRSS2 (HCoV-HKU1) [23]. Notably, HCoV-HKU-1 and HCoV-OC43 are both reported to utilize 9-O acetylated sialic acids (9-O-Ac-Sia) for initial attachment via their spike S1 N-terminal domain (S1A) [24–26], but HCoV-HKU-1 also engages TMPRSS2 for entry [23]. Although no protein receptor is known for HCoV-OC43, entry occurs following endocytosis [27]. Importantly, neither spike 9-O-Ac-Sia interaction, nor low pH/protease treatment triggers conformational changes for membrane fusion [28]. Additionally, while the S1B spike domain is often used for receptor engagement by other coronaviruses, its function in HCoV-OC43 is unknown.

Understanding the mechanisms mediating virus entry can facilitate the design of therapeutics that target virus receptors [29–32].We used genome-wide CRISPR screens and newly established HCoV-OC43 genetics platform to identify host factors and spike sequences driving HCoV-OC43 infection. We identify *NXPE3* as a novel gene required for HCoV-OC43 infection, both *in vitro* and *in vivo*, and show that it is required for 9-O-Ac-Sia biosynthesis. We derived NXPE3/9-O-Ac-Sia-independent HCoV-OC43 derivatives and show that changes in receptor requirement are conferred by amino acid substitutions in either S1A domain or S1B domains that enable heparan sulfate use. These findings reveal surprising flexibility in how HCoV-OC43 spike engages attachment receptors. We further show that interferon-induced transmembrane (IFITM) proteins are required for HCoV-OC43 virion/cell fusion and infection, irrespective of which spike domain glycan drives attachment, indicating a post-attachment fusion-driving role for IFITM proteins in HCoV-OC43 entry.

## Results

### Requirement for NXPE3 in HCoV-OC43 infection

To identify host factors required for HCoV-OC43 (HCoV-OC43(WT), ATCC VR-1588) infection in A549 lung cells, we conducted a genome-wide positive selection CRISPR-survival screen [33]. *NXPE3* was identified as a top-ranked ‘hit’, required for HCoV-OC43(WT) infection **(Fig. 1A, Supplementary table 1)**. To validate this hit, we generated *NXPE3* CRISPR-cas9 knockout cells based on A549-huACE2, 293T cells or primary NHBE cells, using four guide RNAs. *NXPE3* knock-out reduced infectivity in all cell lines, with the largest reduction in 293T cells **(Fig. S1A)**. While the retromer complex component, VPS29, was required for infection by all common cold coronaviruses [33], *NXPE3* requirement was highly specific for HCoV-OC43(WT) **(Fig. S1B)**. Individual single-cell 293T NXPE3-knock out clones **(Fig. S1C)** were nearly completely resistant to HCoV-OC43(WT) infection, but infection was restored by CRISPR-resistant (^re^) NXPE3 expression (NXPE3^re^-HA) **(Fig. 1B)**.

**Fig. 1:**
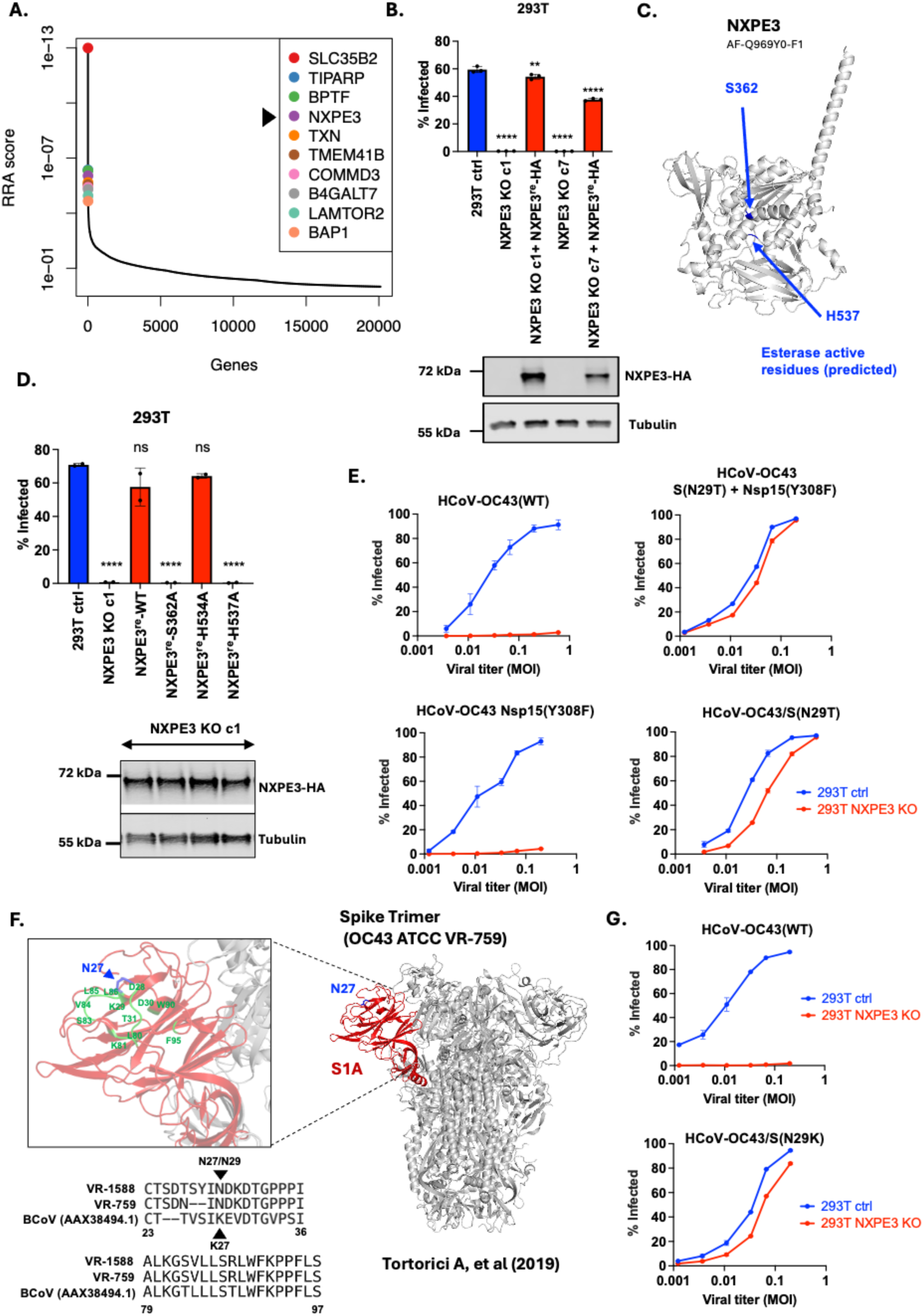
Requirement for NXPE3 in HCoV-OC43 infection. (A) Genome-wide CRISPR-screen robust rank aggregation (RRA) scores for genes required for HCoV-OC43(WT) infection in A549 lung epithelial cells. Top 10 candidates, including NXPE3 are highlighted. n=3 for both infected and uninfected conditions. (B) HEK293T cells (+/-NXPE3 knockout) were infected with HCoV-OC43(WT) for 24 h and the percentage infected cells determined using FACS with antibodies against viral nucleocapsid. NXPE3^re^-HA represents a codon-optimized, guide RNA resistant HA-tagged NXPE3 expression construct. Error bars represent SEM. Each dot indicates a replicate (n=3). P-values are determined by one-way ANOVA (**, p=0.0024; ****, p<0.0001). The lower panel shows a representative immunoblot to assess NXPE3^re^-HA protein levels, with tubulin as a loading control. (C) AlphaFold model of NXPE3 with amino acids predicted to be required for esterase activity marked. (D) HEK293T NXPE3 KO cells expressing various NXPE3^re^-HA mutants were infected with HCoV-OC43(WT) for 24h. The lower panel shows a representative immunoblot to assess NXPE3^re^-HA protein expression with tubulin as a loading control. Replicates are shown as individual dots (n=3) for each experiment accompanied by SEM. P-values are determined by one-way ANOVA (****, p<0.0001; ns: not significant). Infection was assessed by FACS using antibodies against viral nucleocapsid proteins. (E) HEK293T cells (+/-NXPE3 knockout) were infected with HCoV-OC43 viruses with the indicated mutations at various MOIs for 24h. Infection was assessed by flow cytometry using antibodies against viral nucleocapsid proteins. n=2 per MOI, error bars indicate SEM. (F) 3D structure of spike trimer of HCoV-OC43 VR-759 highlighting the spike S1A domain (i.e. S1 NTD) as well as the 9-O-Ac-Sia acid binding site as determined by [28]. The left panel displays amino acid sequence alignment around the 9-O-Ac-Sia acid binding site for HCoV-OC43 ATCC strains as well as a bovine coronavirus isolate (AAX38494.1) which has a N29K substitution. (G) HEK293T cells were infected with the indicated HCoV-OC43 viruses for 24h at various MOIs. Infection was assessed by flow cytometry using antibodies against viral nucleocapsid proteins. n=2 per MOI, error bars indicate SEM.

No function has been reported for NXPE3. The protein is predicted to localize in the secretory pathway and contains a neurexophilin and the Pmr5 and Cas1p (PC)-esterase domain [34]. Alignment of PC-esterase family proteins implicates residues S362 and H537 as esterase active site residues in NXPE3 **(Fig. 1C)** [34]. Unlike wildtype NXPE3^re^-HA, S362A or H537A mutants failed to rescue HCoV-OC43(WT) infection **(Fig. 1D)** suggesting that NXPE3 esterase function was required. We passaged HCoV-OC43(WT) on 293T NXPE3 KO cells **(Fig. S2A)** and after five passages (3-4 weeks), obtained virions bearing nsp15 (Y308F) and spike S1A (N29T) mutations **(Fig. S2B)**. To study these mutations, we built a reverse genetics system (**Fig. S2C, D**) based on the circular polymerase extension reaction (CPER) method [35]. When introduced into the cloned HCoV-OC43, the spike N29T substitution, but not nsp15 Y308F, restored HCoV-OC43 infection in cells lacking NXPE3 **(Fig. 1E)**.

Notably, the HCoV-OC43 spike:9-O-Ac-Sia binding site in S1A includes residue N27 (ATCC VR-759 strain; corresponding to N29 in the ATCC-VR-1588 strain used herein **(Fig. 1F)**) and N-to-A substitution was reported to abolish HCoV-OC43 spike-pseudotype entry [28]. Conversely, a naturally occurring bovine coronavirus isolate encodes an N29K substitution [36] and, like N29T, introducing this change in HCoV-OC43 abolished the NXPE3 requirement **(Fig. 1G)**.

### NXPE3 enables HCoV-OC43 virion binding to cells and is required to generate 9-O-Ac-Sia

Because S1A mutations at the 9-O-Ac-Sia binding conferred NXPE3-independence we reasoned that NXPE3 may be involved in virion binding to cells and/or the generation of 9-O-Ac-Sia. We constructed a virus clone (HCoV-OC43-dns2-M-eGFP) where the accessory gene ns2 was replaced with a second copy of HCoV-OC43(WT) M (membrane) protein, fused to eGFP **(Fig. 2A)**. Incorporation of the M-eGFP fusion protein generates eGFP-labelled virions. We also generated 293T cells lacking CASD1 **(Fig. S3A)**, which attaches acetyl groups to cytidine monophosphate (CMP)-activated sialic acid before conjugation to glycoproteins and lipids [37], or CMAS **(Fig. S3A)** which activates sialic acid by attaching CMP to it [38]. Labelled HCoV-OC43 virions bound to control cells but not to CASD1, CMAS or NXPE3 knockout cells, **(Fig. 2B, C)**, suggesting that NXPE3 is required for HCoV-OC43 virion binding.

**Fig. 2:**
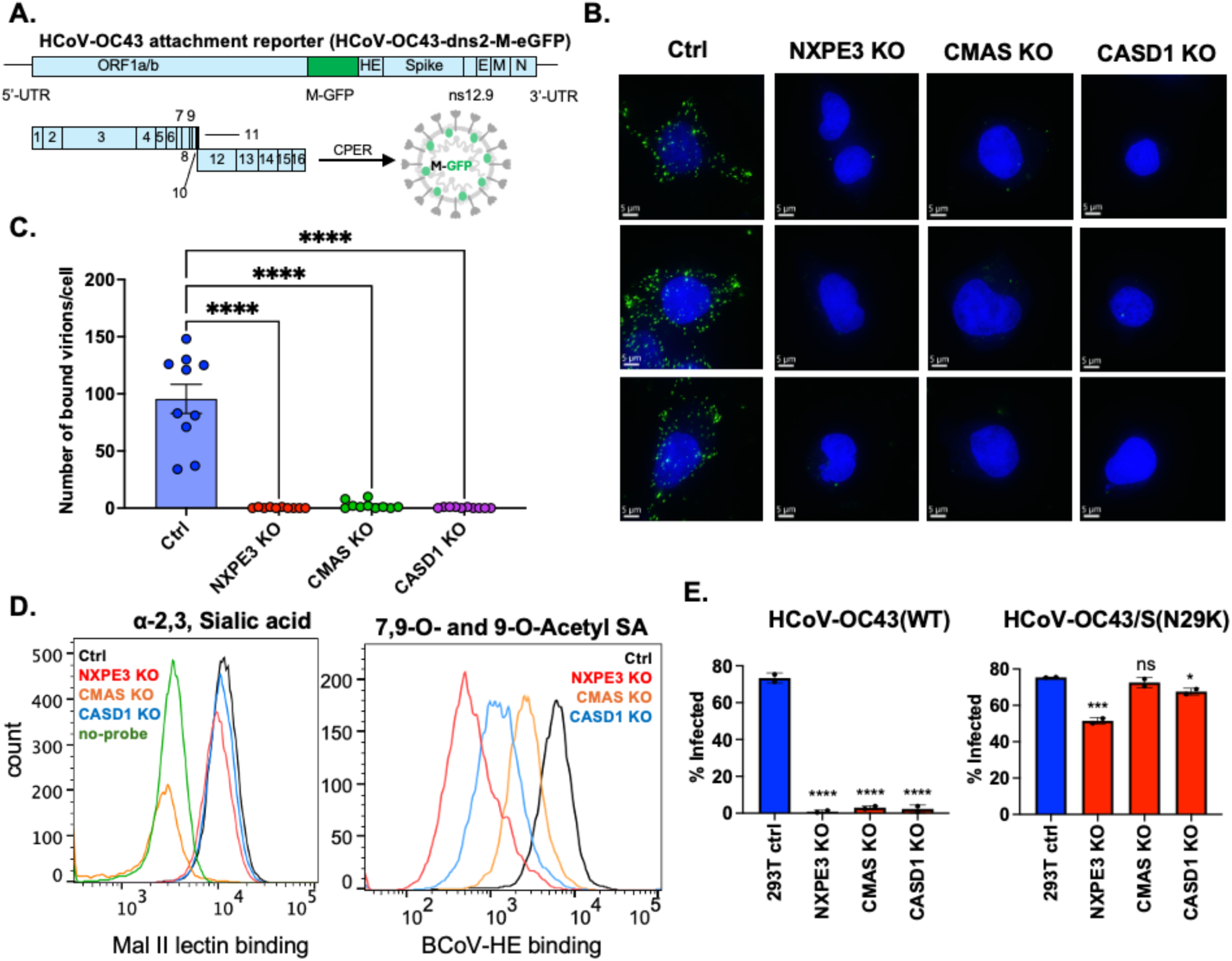
NXPE3 enables HCoV-OC43 virion binding and is required to generate cell surface 9-O-Ac-Sia. (A) Schematic representation of eGFP labelled HCoV-OC43 virions, replacing ns2 with M-tagged eGFP. (B) Virion binding assays were conducted at an MOI:50 on various 293T cells by incubating virions with cells at 4°C for 1.5 hours, followed by fixation and microscopy. Representative images are shown. Hoechst staining (blue) used as nuclear counterstain and virions are shown as green puncta. (C) Graph represents 10 images per condition (from B) obtained from two biological replicates and statistical significance was computed using one-way Anova (****, p<0.0001). Individual cells are shown as individual data point for each condition, bars indicate mean, error bars indicate SEM. (D) Purified BCoV-HE protein from (Fig S3B) was used to bind A549-ACE2 cells to detect bound protein using flow cytometry by staining for His tag while Lectin binding was conducted using MAL II lectin. Representative histograms are shown. (E) HEK293T cells were infected with either HCoV-OC43(WT) or HCoV-OC43/S(N29K) for 24 hours. Infection was assessed by flow cytometry using antibodies against viral nucleocapsid proteins. SEM is shown. Each dot represents a replicate. P-values are determined by one-way ANOVA (****, p<0.0001; ***, p<0.001; **, p<0.005; *, p<0.05; ns: not significant)

The *Embecoviruses*, bovine coronavirus (BCoV), HCoV-OC43, and HCoV-HKU-1 encode a hemagglutinin esterase (HE) protein, that removes cell-surface sialoglycans. We generated an enzymatically inactive Bovine coronavirus HE protein fused to an IgG1 Fc domain and His-tag (HE-Fc-his) as previously described [39] to use as a detection reagent for 7,9-O- and 9-O-acetylated sialic acids **(Fig. S3B)**. Flow cytometric binding assays on A549-huACE2 cells lacking NXPE3, CMAS, CASD1 **(Fig. 2D)** revealed little or no binding of BCoV HE-Fc-His protein to cells lacking NXPE3, Conversely, MAL II lectin binding to α-2,3 sialic acids [40] was sensitive only to CMAS KO **(Fig. 2D)**. Finally, 293T cells lacking NXPE3, CASD1, or CMAS supported HCoV-OC43/S(N29K) infection, but not HCoV-OC43(WT) infection **(Fig. 2E)**. We conclude that NXPE3 is required for 9-O-Ac-Sia synthesis, but that an S1A substitution (N29K) confers 9-O-Ac-Sia independence.

### NXPE3 expression determines susceptibility to HCoV-OC43 *in vitro* and in *vivo*

Through inspection of gene expression databases, we identified the hepatoma cell line, HuH1 as lacking NXPE3 expression and confirmed this finding using an RT-PCR assay **(Fig. 3A)**. Accordingly, HuH1 cells were not susceptible to infection by HCoV-OC43(WT) but could be infected by HCoV-OC43/S(N29K) (**Fig. 3B)**. Susceptibility of HuH1 cells to HCoV-OC43(WT) was conferred by NXPE3^re^-HA expression but not by NXPE3^re-^HA (S362A/H537A/H534A), consistent with results in 293T NXPE3 KO cells **(Fig. 3C)**. To investigate whether NXPE3 was required for *in vivo* HCoV-OC43 infection, we used a mouse model [3]. Note that cell-culture adapted HCoV-OC43 is usually avirulent in adult C57BL/6 mice, while a neurovirulent HCoV-OC43 strain (HCoV-OC43_nv_) derived from suckling mouse brains following inoculation with material directly obtained from human cases causes lethal infection in adult mice [3]. We generated C57BL/6 mice lacking NXPE3 (NXPE3-/-) whose splenocytes display poor binding to the BCoV-HE-Fc-His protein, in line with our characterization of NXPE3 **(Fig. 3D)**. We infected both C57BL/6 and NXPE3-/- mice with a lethal dose of HCoV-OC43_nv_ and harvested RNA from mouse brain homogenates at 5 days postinfection. HCoV-OC43_nv_ was lethal to wildtype mice, with 9/10 mice succumbing to infection by 10dpi while all 10 NXPE3-/- mice remained healthy **(Fig. 3E)**. Moreover, while HCoV-OC43_nv_ viral RNA was readily detected in infected C57BL/6 mice brains, no viral RNA was detected in infected NXPE3-/- mice brains **(Fig. 3F)**. NXPE3, is therefore required for HCoV-OC43 susceptibility and pathogenesis in mice.

**Fig. 3:**
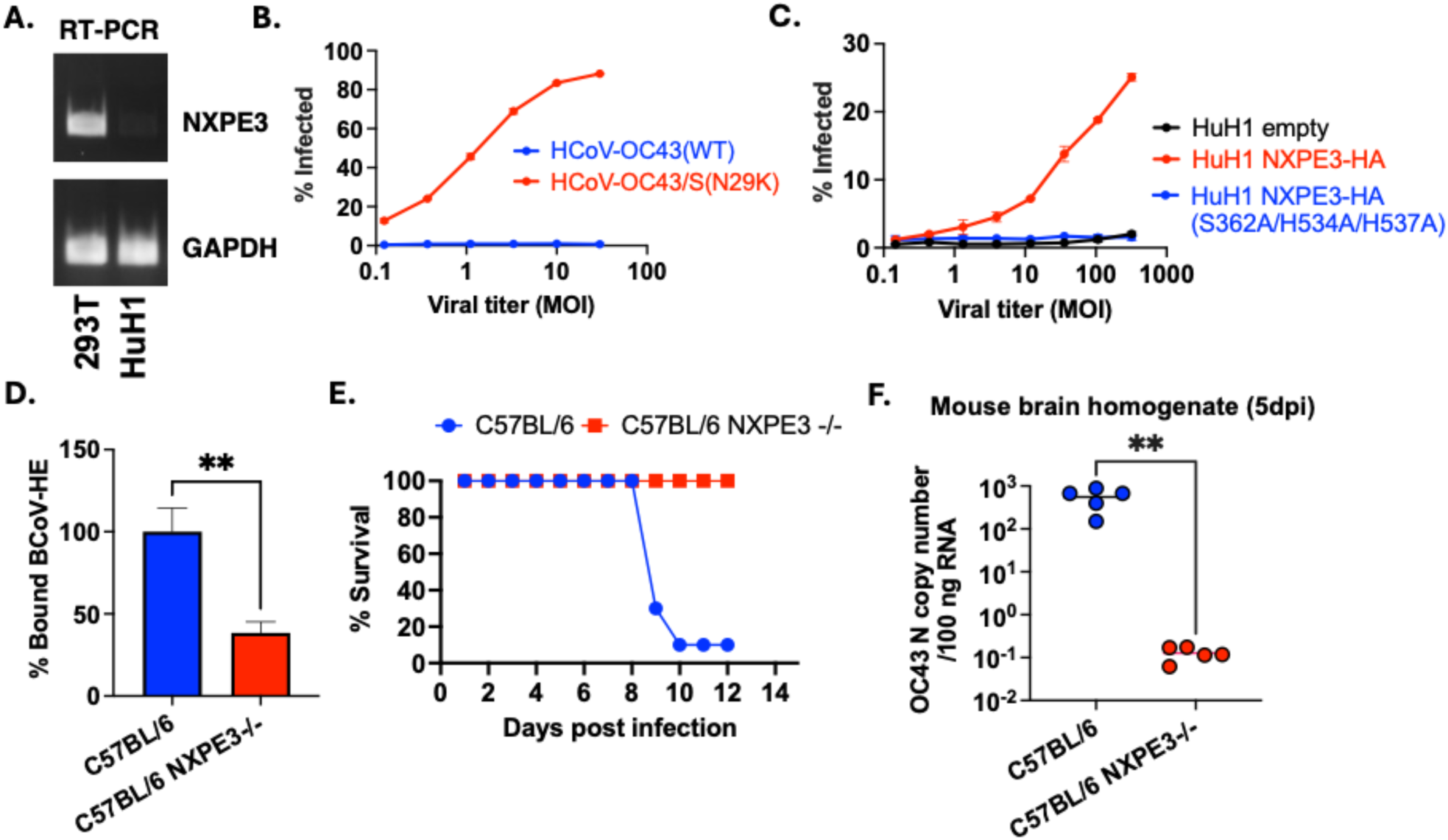
NXPE3 determines susceptibility to HCoV-OC43 *in vitro* and in *vivo*. (A) RT-PCR detection of NXPE3 mRNA in Huh-1 and 293T cells as visualized by agarose gel electrophoresis. (B) Huh1 cells infected with HCoV-OC43(WT) and HCoV-OC43/S(N29K) for 24 h. Infection was assessed by flow cytometry using antibodies against viral nucleocapsid protein (n=3 per MOI, error bars represent SEM). (C) HCoV-OC43(WT) infection assessed on various HuH1-based cell lines. (n=2 per MOI, error bars represent SEM) (D) Splenocytes isolated from C57BL/6 mice were resuspended in RPMI medium, filtered and fixed to assess binding of BCoV-HE protein to cells, as in Fig. 2D. Normalized binding is shown for 3 spleens per mice group. Student’s t-test was performed (**, p=0.0092), error bars represent SEM. (E) C57BL/6 mice (+/-NXPE3 KO) were intranasally infected with HCoV-OC43_nv_ and followed to assess survival over time (ten mice per group). (F) C57BL/6 mice (+/-NXPE3 KO) were intranasally infected with HCoV-OC43_nv_. At 5dpi, mice brains were extracted and homogenized in 1X PBS, followed by RNA extraction and qPCR to measure viral genome copy numbers. Each dot represents a biological replicate. Student’s t-test was performed (**, p=0.0024).

### HCoV-OC43/S(N29K) infection depends on heparan sulfate

The NXPE3/9-O-Ac-Sia-independent HCoV-OC43 mutants S(N29T and N29K) presumably engage alternative attachment receptors. To identify these, we challenged 293T NXPE3 KO cells with HCoV-OC43/S(N29K), following transduction with vectors carrying sgRNAs that target transmembrane proteins, selected based on their ranking in our genome-wide screen **(Fig. 1A)**. Notably, *B4GALT7* (Beta-1,4-galactosyltransferase 7), an enzyme in the glycosaminoglycan biosynthesis pathway [41], emerged as the only hit **(Fig. 4A)**. Glycosaminoglycans such as heparan sulfate (HS) are used by several viruses for attachment [42, 43]. We found that 293T cells engineered to lack B4GALT7 **(Fig. S4A)** were resistant to HCoV-OC43/S(N29K) infection and susceptibility could be restored by expression of B4GALT7^re^-HA **(Fig. 4B)**. In contrast, 293T B4GALT7 KO cells remained susceptible to HCoV-OC43(WT) **(Fig. 4C)**. Soluble heparin inhibited HCoV-OC43/S(N29K) infection of 292T cells (IC_50_=0.13156 IU/ml) while HCoV-OC43(WT) was comparatively resistant (IC_50_=3.118 IU/ml) (**Fig. 4D)**. Similar to 293T cells, HCoV-OC43(WT) infected both CHO-K1 cells and a derivative, pgsA745, that lacks glycosaminoglycans [44]. Conversely, CHO-K1 cells were infected by HCoV-OC43/S(N29K) while pgsA745 cells were resistant **(Fig. 4E)**.

**Fig. 4:**
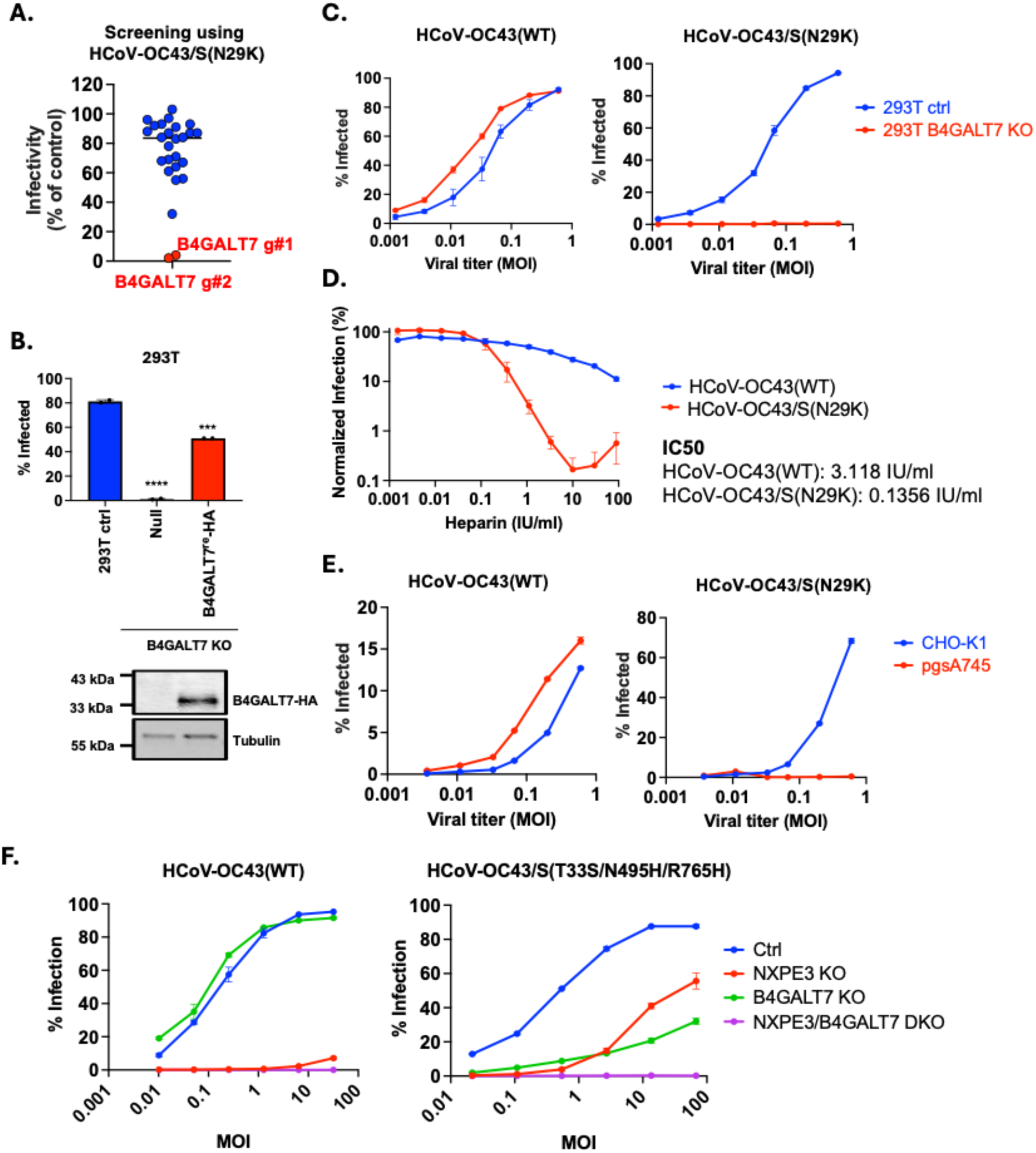
HCoV-OC43/S(N29K) infection depends on heparan sulfate. (A) Small-scale CRISPR screen conducted in 293T cells focused on transmembrane protein hits (from screen in Fig. 1A), assessing HCoV-OC43/S(N29K) infectivity in polyclonal CRISPR-cas9-edited cell lines. Infectivity is normalized to control gRNA treatment. (B) 293T cells infected with HCoV-OC43/S(N29K) for 24 h, infection assessed by flow cytometry using antibodies against viral nucleocapsid protein. B4GALT7^re^-HA represents a codon-optimized guide RNA-resistant HA-tagged B4GALT7 (n=2, bar represents SEM). P-values are determined by one-way ANOVA (****, p<0.0001; ***, p<0.001). The lower panel shows an immunoblot to assess B4GALT7^re^-HA protein levels with tubulin as a loading control. (C) HEK293T cells were infected with either HCoV-OC43(WT) or HCoV-OC43/S(N29K) for 24 hours at various MOIs, infection assessed by flow cytometry using antibodies against viral nucleocapsid protein (n=3 per MOI, error bars represent SEM). (D) HEK293T cells were incubated with either HCoV-OC43(WT) or HCoV-OC43/S(N29K) (MOI:0.5) for 24 h with various concentrations of heparin (IC_50_ concentrations shown, n=3 per treatment condition, error bars represent SEM). (E) CHO cell derivatives infected with either HCoV-OC43(WT) or HCoV-OC43/S(N29K) for 24 h. (n=2 per MOI, error bars represent SEM). Infection was assessed by flow cytometry using antibodies against viral nucleocapsid protein. (F) HEK293T-based cell lines infected with either HCoV-OC43(WT) or HCoV-OC43/S(T33S/N495H/R765H) for 24 h (n=2 per MOI, error bars represent SEM). Infection was assessed by flow cytometry using antibodies against viral nucleocapsid protein.

Unlike our findings in 293T cells, HCoV-OC43(WT) infection was previously reported to be partly dependent on glycosaminoglycan synthesis in A549 and Huh7 cells (without a requirement for N29K mutation) [45, 46]. Because 293T cells express low levels of HS [47], we hypothesized that HCoV-OC43(WT) can in fact use either 9-O-Ac-Sia or HS as attachment receptors but shows greater dependence on 9-Ac-O-Sia in 293T cells. HS is displayed on the surface of cells as proteoglycans (HSPG), conjugated to syndecan or glypican protein cores. We analyzed publicly available RNA-seq data across cell lines and found that syndecan and glypican proteins are poorly expressed in 293T cells **(Fig. S4B)**. Overexpression of syndecan-2 in 293T NXPE3 knockout cells both increased cell surface HS, and restored susceptibility to HCoV-OC43(WT) **(Fig. S4C, S4D)**. Hence, the ability of HCoV-OC43 to utilize HSPG depends on both viral spike and cellular HSPG determinants. Moreover, HCoV-OC43/S(N29T) could apparently use both 9-O-Ac-Sia and HSPG, as knockout of both NXPE3 and B4GALT7 was required to abrogate HCoV-OC43/S(N29T) infection **(Fig. S4E)**. Interestingly, addition of three substitutions (S33/H495/H765) to HCoV-OC43(WT), which were acquired by HCoV-OC43_nv_ *in vivo* [3], also conferred ability to use HSPG (**Fig. 4F)**, while retaining 9-O-Ac-Sia usage.

### Acquisition of substitutions in a different HCoV-OC43 spike domain (S1B) can confer HSPG usage

Substitutions in S1A (at N29) conferred changes in the use of 9-O-Ac-Sia or HSPG. We passaged HCoV-OC43/S(N29T) on 293T NXPE3/B4GALT7 double-KO (DKO) cells, which we expected to lack both 9-Ac-O-Sia and HS attachment factors. After 6 passages, A virus that encoded two additional substitutions at spike (N514K/D535N) and infected 293T NXPE3/B4GALT7 DKO cells emerged **(Fig. 5A, B)**. These residues are within the S1B domain, which heretofore had no defined function in HCoV-OC43 entry **(Fig. 5B)** [48]. To determine if these mutations altered receptor usage, we conducted another genome-wide CRISPR screen, in 293T NXPE3/B4GALT7 DKO target cells, selecting those that survived HCoV-OC43/S(N29T/N514K/D535N) infection. Two genes, *IFITM1* and *EXT2* emerged as top ranked hits **(Fig. 5C)**. The *EXT2* hit was surprising, as it encodes an enzyme that (like B4GALT7) participates in glycosaminoglycan synthesis [49], which was expected to be absent in 293T NXPE3/B4GALT7 DKO cells. That *EXT2* emerged as a hit suggested that *B4GALT7* knockout may not completely eliminate glycosaminoglycans, and that HCoV-OC43/S(N29T/N514K/D535N) was able to use trace residual HSPGs remaining on 293T NXPE3/B4GALT7 DKO cells. Thus, we generated 293T NXPE3/B4GALT7/EXT2 triple-knockout (TKO) cells and confirmed that EXT2 was important for HCoV-OC43/S(N29T/N514K/D535N) infection **(Fig. 5D** and **S5)**. To further determine whether the S1B substitutions (N514K/D535N) facilitated HSPG usage, we constructed a HCoV-OC43/S(N514K/D535N), retaining wildtype N29. HCoV-OC43/S(N514K/D535N) infected 293T NXPE3/B4GALT7 DKO cells, showing that the S1B substitutions alone could enable infection of cells with trace HSPGs **(Fig. 5E)**. Notably, HCoV-OC43/S(N514K/D535N) was highly sensitive to heparin (IC_50_=0.00052 IU/ml), consistent with the notion that HCoV-OC43/S(N514K/D535N) S1B could use HS **(Fig. 5F)** for entry.

**Fig. 5:**
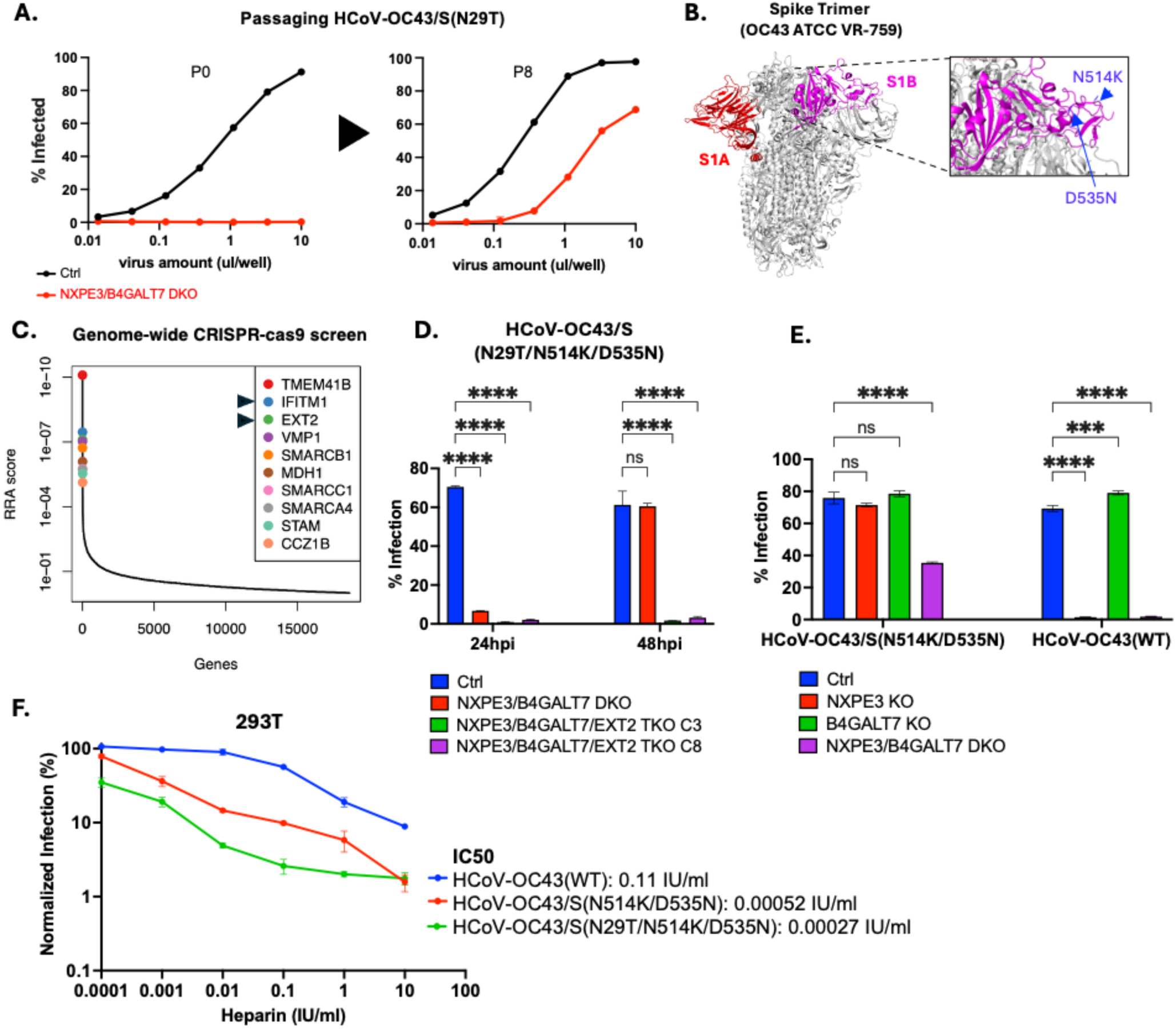
Acquisition of substitutions in the HCoV-OC43 S1B spike domain that confer HSPG usage. (A) Passaging of HCoV-OC43/S(N29T) on 293T NXPE3/B4GALT7 double KO (DKO) cells generates NXPE3 and B4GALT7-independent HCoV-OC43. Infectivity of supernatant collected from passage 0 (P0) and passage 8 (P8) is shown using various virus volumes (n=1 per MOI). (B) Substitutions N514K and D535N in the S1B spike domain. (C) Robust rank aggregation scores for genes required for HCoV-OC43/S(N29T/N514K/D535N) infection in NXPE3/B4GALT7 DKO 293T cells (analysis similar to Fig. 1A). Top 10 genes including *IFITM1* and *EXT2* are highlighted. N=3 for both infected and uninfected conditions. (D) HEK293T cells infected with HCoV-OC43/S(N29T/N514K/D535N) at an MOI:1 for 48 hours. Infection was assessed by flow cytometry using antibodies against viral nucleocapsid proteins (n=3, error bars represent SEM). P-values are determined by Two-way ANOVA (****, p<0.0001; ns: not significant). (E) 293T cell derivatives infected with HCoV-OC43/S(N514K/D535N) or HCoV-OC43(WT) at an MOI:1 for 24 hours. Infection was assessed by flow cytometry using antibodies against viral nucleocapsid proteins (n=3, error bars represent SEM). P-values are determined by Two-way ANOVA (****, p<0.0001; ***, p<0.001; ns: not significant). (F) Heparin sensitivity experiment was conducted as in Fig. 4D for different viruses (MOI:0.2). (n=2 for each concentration of heparin; IC_50_ values shown). Error bars indicate SEM. Infection was assessed by flow cytometry using antibodies against viral nucleocapsid proteins.

### IFITM proteins mediate HCoV-OC43 virion-cell fusion, irrespective of virion attachment mechanism

IFITM1/2/3 proteins are interferon-inducible type II transmembrane endo/lysosomal proteins that hinder entry of enveloped viruses such as IAV [50], HIV-1 [51] Paradoxically, IFITM2/3 proteins have been reported to enhance HCoV-OC43 susceptibility [52]. While IFITM1 emerged as a top-ranked hit in our screen, the gRNAs targeting IFITM1 and IFITM3 exhibit high sequence similarity, and such gRNAs may have acted through both targets. We generated cell lines that lacked IFITM1/2/3, using a sgRNA that targets both IFITM 2 and 3 **(Fig. S6A, B)**. As the IFITM1 locus is flanked by IFITM2 and IFITM3, CRISPR-mediated DNA breaks in IFITM2 and IFITM3 could functionally delete all three genes **(Fig. S6A).** Absence of IFITM1/2/3 impaired HCoV-OC43 infection irrespective of the aforementioned spike mutations that changed attachment factor dependence **(Fig. 6A).** Infection of 293T IFITM1/2/3 knockout cells was restored by gRNA-resistant (^re^) IFITM2^re^ or IFITM3^re^, and to some extent by IFITM1. Notably, IFITM2^re^ or IFITM3^re^ derivatives with mutated endocytic signals[53, 54] which changes localization to the plasma membrane, (IFITM2Y19A^re^ and IFITM3Y20A^re^), also rescued HCoV-OC43 infection **(Fig. S6C),** suggesting that IFITM endocytic trafficking is not required for HCoV-OC43 infection. Both the number and size of infected foci in focus formation assays was increased by IFITM1/2/3 further confirming their effect on HCoV-OC43 infection **(Fig. S6D-F)**.

**Fig. 6:**
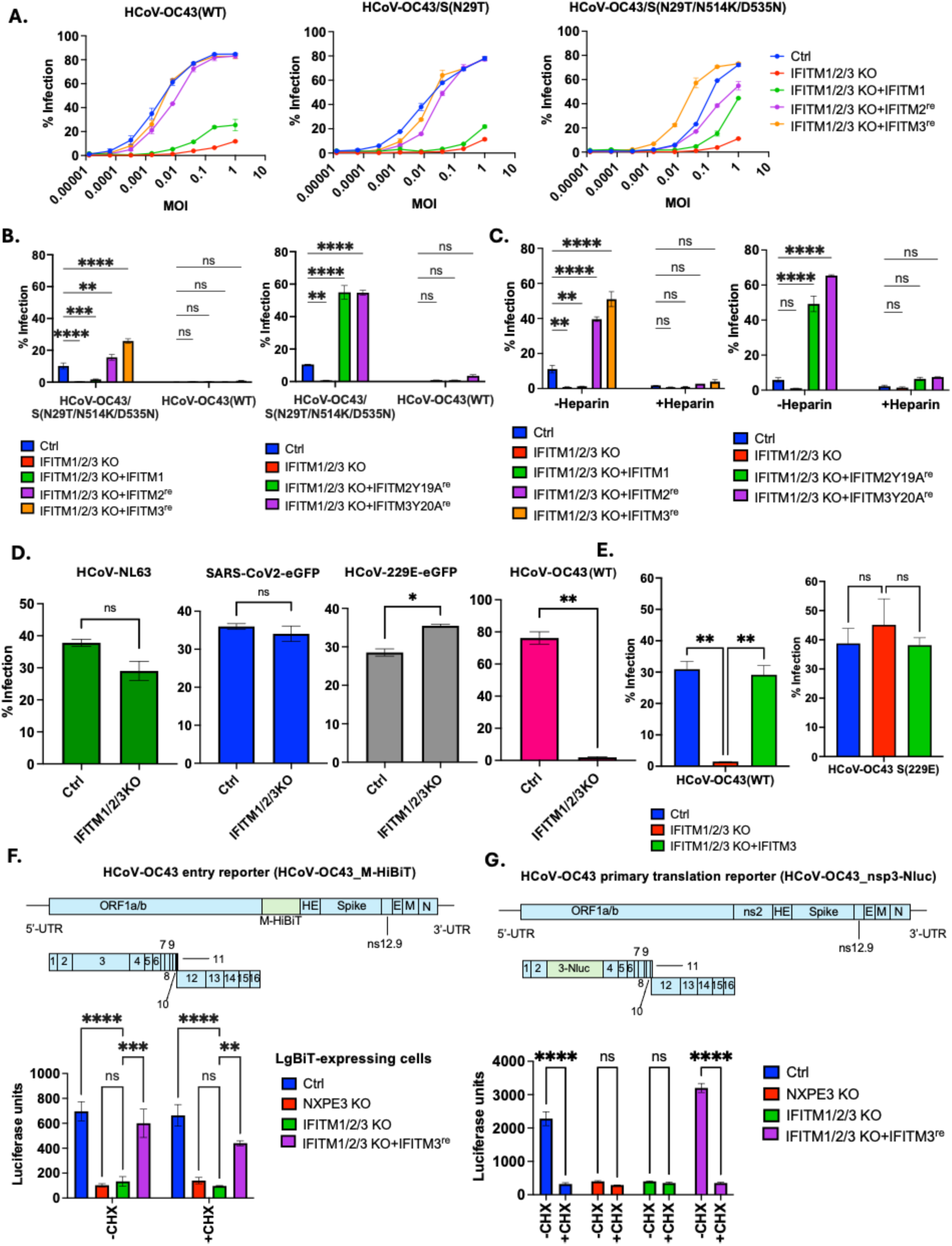
IFITM proteins are required for HCoV-OC43 virion-cell fusion, irrespective of the attachment factor. (A) HCoV-OC43 infection in unmanipulated 293T, and IFITM1/2/3 knockout cells with IFITM1, 2 or 3, expressed using guide RNA resistant codon-optimized IFITMs(^re^) at 24 h (n=2 per MOI, SEM is shown). Infectivity is determined by flow cytometry using antibodies against viral nucleocapsid proteins. (B) NXPE3/B4GALT7 DKO 293T-based cell lines infected with HCoV-OC43(WT) or HCoV-OC43/S(N29T/N514K/D535N) viruses for 48 h to assess infectivity (n=2, SEM is shown). Infection was assessed by flow cytometry using antibodies against viral nucleocapsid proteins. P-values are determined by Two-way ANOVA (****, p<0.0001; ***, p<0.001; **, p<0.005; ns: not significant). (C) NXPE3/B4GALT7 DKO 293T-based cell lines were treated with heparin (10 IU/ml) and simultaneously infected with HCoV-OC43/S(N29T/N514K/D535N) for 48 h to assess infectivity. Infection was assessed by flow cytometry using antibodies against viral nucleocapsid proteins (n=2, error bars represent SEM). P-values are determined by Two-way ANOVA (****, p<0.0001; **, p<0.005; ns: not significant). (D) HCoV-OC43(WT) was used to infect 293T-based cell lines and 293T-based cell lines expressing ACE2 or hAPN were used for infection with SARS-CoV2-dORF7-eGFP/ HCoV-NL63 or HCoV-229E-eGFP as appropriate for 24 hours (n=3 for each virus, error bars represent SEM). Infection was assessed by flow cytometry using antibodies against viral nucleocapsid proteins or using eGFP reporter expression. Student’s t-test was performed (*, p=0.0121; **, p<0.0047; ns: not significant). (E) Cell lines (293T-based) transduced with hAPN were infected with either HCoV-OC43(WT) or HCoV-OC43 S(229E) for 24 hours followed by immunostaining for nucleocapsid protein and flow cytometry. Infectivity is shown for n=3 (error bars represent SEM). one-way Anova was conducted (**, p<0.005; ns: not significant). Live-cell NanoLuc luciferase assays were conducted for HCoV-OC43 fusion reporter (F) and primary translation reporter (G) at an MOI=10 and 2 respectively. Luciferase values are plotted for n=3 with SEM at 2hpi (F) and 4hpi (G). Statistical significance was computed with two-way Anova. (****, p<0.0001; ***, p<0.001; **, p<0.005; ns: not significant)

In cells depleted of attachment factors (293T NXPE3/B4GALT7 KO) – with residual trace amounts of glycosaminoglycans, IFITMs were required for HCoV-OC43/S(N29T/N514K/D535N) infection **(Fig. 6B)**. Notably, in this context, overexpression of IFITM2^re^ or IFITM3^re^ (or particularly IFITM2Y19A^re^ and IFITM3Y20A^re^ **(Fig. 6B)**) markedly increased infection compared to unmanipulated 293T NXPE3/B4GALT7 KO cells, suggesting that IFITM2^re^/3^re^ overexpression enhances infection when attachment is sub-optimal. However, IFITM1/2/3 overexpression did not rescue HCoV-OC43(WT) infection of 293T NXPE3/B4GALT7/IFITM1/2/3 KO cells **(Fig. 6B)**, suggesting that a glycan attachment factor is required even in the context of IFITM overexpression. Additionally, HCoV-OC43/S(N29T/N514K/D535N) infection of 293T NXPE3/B4GALT7/IFITM1/2/3 KO cells overexpressing IFITMs was inhibited by heparin, also consistent with an ongoing requirement for glycan mediated attachment in the context of IFITM overexpression **(Fig. 6C)**.

Notably, infection by other coronaviruses tested (HCoV-NL63, HCoV-229E and SARS-CoV-2) in 293T cells engineered to express the appropriate receptors (ACE2 and hAPN**, Fig. S6G**) was unaffected by IFITM1/2/3 **(Fig. 6D)**. Infection by a chimeric HCoV-OC43 whose spike was replaced by that of HCoV-229E spike was also unaffected by IFITM3^re^, showing that IFITM3^re^ effects are HCoV-OC43 spike-dependent (**Fig. 6E**). To gain mechanistic insights, we used two HCoV-OC43 indicator viruses [55]. The first was based reconstitution of split NanoLuc luciferase comprising two fragments (HiBiT and LgBiT). First, LgBiT was expressed in the cytosol of target cells. Then the non-essential HCoV-OC43 ns2 protein was replaced with a second copy of the M protein fused to HiBiT, resulting in HiBiT incorporation into virions. Thus, virion-cell fusion (entry) generates luminescence through (HiBiT/LgBiT) reconstitution **(Fig. 6F, S6H)**. A second indicator virus incorporated a NanoLuc luciferase orf, fused to the N-terminus of NSP3, which generates luminescence in target cells upon translation of the incoming viral genome **(Fig. 6G)**. Using the HCoV-OC43_M-HiBiT reporter, the luminescence signal resulting from virus-cell fusion was reduced by ∼6-fold by NXPE3 or IFITM1/2/3 knockout and restored by IFITM3^re^ in the latter **(Fig. 6F).** The signal was not due to new M-HiBiT protein synthesis, as evidenced by its resistance to cycloheximide (CHX). The luminescence signal resulting from translation of the incoming viral RNA 4h after HCoV-OC43_nsp3-Nluc infection was CHX-sensitive, similarly reduced ∼8-fold by NXPE3 or IFITM1/2/3 KO, and restored in IFITM1/2/3 KO cells by IFITM3^re^ expression **(Fig. 6G)**. Overexpressing IFITM3Y20A^re^ did not rescue virion binding that had been ablated by NXPE3/B4GALT7 KO **(Fig. S7A, B)** and 293T cells lacking IFITM1/2/3 (but were otherwise unmanipulated) bound virions similarly to parental cells **(Fig. S7C, D)**.

Overall, these data indicate that IFITMs act downstream of attachment to drive HCoV-OC43 virion-cell fusion and complete the viral entry process. Consistent with this model, analysis of IFITM3 mutants **(Fig. S8A-C)** showed that membrane-associated portions, including the amphipathic helices (AH1 & AH2) and palmitoylation sites (C71/72) were necessary for infection, while deletions in the N-terminal cytosolic (Δ21) or C-terminal extracellular tails (Δ2 and Δ4) did not affect activity **(Fig. S8B)**. IFITM1 was a comparatively poor enhancer of HCoV-OC43 infection, but a C-terminal tail truncation increased HCoV-OC43 infection (IFITM1 C-term del) **(Fig. S8B)**, Thus, the extracellular C-termini of IFITM proteins can influence potency but are dispensable for HCoV-OC43 infection.

## Discussion

We identify NXPE3 as essential for the synthesis of 9-O-Ac-Sia, a critical HCoV-OC43 attachment receptor, both *in vitro* and *in vivo*. The sialic acid esterase (SIAE) is the only esterase known to participate in sialic acid biosynthesis, removing acetyl groups at C-4/C-9 positions of sialic acid [56]. NXPE3 likely functions analogously to SIAE but on different intermediates, thus generating differentially acetylated sialic acids. Notably, knockout of NXPE3 yielded apparently healthy mice, that were resistant to lethal challenge with HCoV-OC43_nv_. Whether NXPE3 is required by other viruses that use 9-O-Ac-Sia such as influenza C is not known [57]. Our findings raise the possibility that sialic acid modifying enzymes like NXPE3 might be therapeutically targeted to combat virus infections.

Adaptative (N29T) or naturally occurring (N29K) [36] substitutions in S1A conferred NXPE3/9-O-Ac-Sia independence. HCoV-OC43/S(N29K) apparently switched to HS-dependent entry at the expense of 9-O-Ac-Sia utilization and required B4GALT7, a galactosyltransferase that links HS (heparan sulfate) to cell-surface syndecan/glypican proteins [41]. Conversely, HCoV-OC43/S(N29T) virus could utilize both 9-O-Ac-Sia and HS. While HS-dependent entry by HCoV-OC43 was previously reported [45, 46], the ability of HCoV-OC43 to use HS depends on both the spike protein and cellular HSPG expression - the S(N29K/T) substitutions enabled HCoV-OC43 to utilize the low levels of HS normally present on 293T cells [47] while syndecan-2 overexpression [58] allowed HCoV-OC43(WT) to utilize HS for entry therein. Thus, HCoV-OC43 spike variants, exhibit differential propensity to use carbohydrate attachment receptors. Interestingly, S1A in clinical isolates exhibits variability in carbohydrate binding specificity [59]; HCoV-OC43 laboratory strains and HKU-1 coronavirus bind to both 9-O-Ac, 5-N-Ac groups and the 9-*O*-Ac- *α* -2–8-linked-Sia moiety of the 9-O-Ac-GD3_ap_ ganglioside, but HCoV-OC43 clinical isolates bind only the latter. Interestingly, the HCoV-OC43_nv_ strain which has never been cultured in cell lines, was strictly dependent on NXPE3 in mice, but placement of its defining mutations into HCoV-OC43(WT) enabled dual 9-O-Ac-Sia and HS usage. How HCoV-OC43 spike S1A mutations that occur naturally shape glycan usage warrants further investigation.

Adaptive substitutions (N514K/D535N) in a distinct domain of HCoV-OC43 spike (S1B) that enabled HCoV-OC43 to replicate in NXPE3/B4GALT7-DKO cells conferred a dependence on EXT2. This glycosyltransferase that adds heparan sulfate chains to cell-surface proteins [49, 60]. and was expected to be redundant with B4GALT7 in terms its requirement by HCoV-OC43. Our finding suggests plasticity in galactosyltransferase behavior. Moreover, that HCoV-OC43/S(N29T/N514K/D535N) infection was abolished in 293T NXPE3/B4GALT7/EXT2 TKO cells, and that S1B substitutions conferred high sensitivity heparin, indicates HS usage by these S1B substituents. While a role for S1B in HCoV-OC43 entry has not previously been reported, S1B can be targeted by HCoV-OC43 neutralizing antibodies[48]. Our results suggest a model in which HCoV-OC43 infection is initiated by engaging attachment factors via either S1A or S1B domains, with striking naturally occurring plasticity in HCoV-OC43 attachment factor use.

A previous report indicated that the antiviral interferon-induced IFITMs are, paradoxically, required for HCoV-OC43 infection [52], The IFITM proteins emerged in our CRISPR screens as required for HCoV-OC43 infection, but were not required by other coronaviruses. Indeed, we found that replacement of the HCoV-OC43 spike with that of HCoV-229E abolished IFITM1/2/3-dependence. By inclusion of reporter proteins (GFP/ HiBiT) or translatable reporter mRNA (NanoLuc luciferase) into HCoV-OC43 particles, we found that IFITM1/2/3 have no effect on virion attachment but act subsequently to mediate virion-cell membrane fusion. Entry mediated by IFITM3 was dependent on the integrity of its intra-membrane domains, including the palmitoylation sites and amphipathic helices. Although IFITM proteins exert antiviral effects by inhibiting membrane fusion [61], HCoV-OC43 may engage IFITM proteins, but with the opposite consequence, to facilitate entry. While further studies are necessary to elucidate the details of IFTM function in viral entry, this study highlights how HCoV-OC43 exhibits great flexibility in its use of glycan receptors for attachment, enabling the subsequent hijacking of IFITM proteins for virion-cell fusion.

## Online Methods

### Cell culture

A549, 293T (HEK293T), HuH7, Huh1, VeroE6, NHBE and engineered derivatives were grown in DMEM media (Gibco # 11995065) supplemented with 10% fetal-calf serum (Sigma #F0926) and 50 µg/ml Gentamicin solution (Gibco # 15750060). CHO-K1 and pgsA745 cells were cultured in Ham’s F-12 Nutrient Mix medium (Gibco #11765054) with 10% fetal-calf serum and 50 µg/ml Gentamicin solution. Expi293F (ThermoFisher, #A14635) were grown in Expi293 Expression Medium (Gibco # A1435101) under shaking conditions in 125 mL Biodegradable Erlenmeyer Flasks. All cells were maintained at 37°C at 5% CO_2_. Cells infected with authentic CPER-generated coronaviruses were incubated at 34°C at 5% CO_2_ to mimic physiological temperatures of the upper respiratory pathway. All cell lines genetically modified to knockout or overexpress proteins are listed in the Supplementary Table 2. Selection of cell lines were conducted at the following concentrations: 2.5 µg/ml Puromycin, 5 µg/ml Blasticidin, 50 µg/ml Hygromycin B, 100 µg/ml Zeocin and 100 µg/ml G418.

### Virus stock production

All coronavirus stocks were grown in BSL-2 (HCoV-OC43, HCoV-229E-eGFP and HCoV-NL63) or BSL-2+ conditions (SARS-CoV2). SARS-CoV2-dORF7a/b-eGFP virus was generated using a CPER-based reverse genetic system [35] to ensure no BSL-3 grown isolates were used throughout the study. Crude virus supernatants generated by CPER (see method details) were used to infect 5-layer flasks of 293T cells (HCoV-OC43 viruses), VeroE6 cells (HCoV-NL63, SARS-CoV2-eGFP) or HuH7 (wildtype or HCoV-229E-eGFP) at 34°C and 5% CO_2_ conditions. Supernatants were harvested after obtaining 90% cytopathic effects (cell rounding and detachment) and filtered (0.2 micron). Filtered virus stock was then subject to 20% sucrose cushion ultracentrifugation. Virus pellets were resuspended in serum-free DMEM to generate purified virus aliquots. Viruses were titrated by focus formation assays on 293T (HCoV-OC43), 293T-hAPN (HCoV-229E) or 293T-ACE2 cells (HCoV-NL63 and SARS-CoV2) using a 1% CMC overlay. Foci were visualized by eGFP expression or immunostaining against viral nucleocapsid proteins at 4-5 dpi. Information on all viruses used can be found in the Supplementary Table 2. For HCoV-OC43_nv_, two- to three-day-old suckling mice were intracranially infected with 5 x 105 SMLD50 (Suckling Mouse LD50) of HCoV-OC43_nv_. At 2 days post-infection, the mice were decapitated. Heads were pooled and homogenized in sterile PBS, after which the supernatant was clarified by centrifugation. Finally, the total clarified supernatant was aliquoted into 50 uL volumes and stored at -80°C until further use. All procedures used were approved by the University of Iowa Institutional Animal Care and Use Committee and met the stipulations of the Guide for the Care and Use of Laboratory Animals (National Academies Press, 2011).

### Mouse infections and RT-qPCR

Five-eight-week-old C57BL/6 mice were purchased from Jackson Laboratory (#000664) and housed in a standard BSL1 facility at the Rockefeller Comparative Biosciences Center and regular chow diet, water *ad libitum* under a 12-h light–dark cycle. All mice were shifted to BSL2, and experiments were conducted at least 48-72 hours later. The mouse studies were performed in compliance with the animal protocol 24016-H and the Rockefeller University Institutional Animal Care and Use Committee (IACUC). C57BL/6 ýNXPE3 mice were generated by CRISPR & Genome Editing Resource Center (Transgenic & Reproductive Technology Resource Center) at the Rockefeller University using a gRNA targeting mouse NXPE3 at Exon 5 (4^th^ coding exon): taatgaccccgacaacatta (PAM: cgg). NXPE3 -/- mice were genotyped by PCR and Sanger sequencing across the NXPE3 locus using genomic DNA from mouse tail isolated by DNeasy Blood & Tissue kit (Qiagen #69504) according to manufacturer. C57BL/6 or C57BL/6 NXPE3 -/- mice were infected intranasally (1:4 dilution of virus stock, 1:3 lethal dose for survival experiments). Mouse brains were harvested at 5dpi and homogenized in 1X PBS followed by Trizol-based RNA extraction based on manufacturers’ protocol. One-step cDNA and qPCR was performed using Power SYBR Green RNA-to-C_T_ 1-Step kit (Applied biosystems # 4389986) as per manufacturers’ conditions. Virus titers were determined by comparing C_T_ values with a purified and titrated virus stock of ATCC-1588 HCoV-OC43(WT). For survival experiments, mice were simply monitored to assess weight and lethality. All experiments were conducted using mixed populations of males and females. Primers used for qPCR and genotyping are listed in the Supplementary Table 2. qPCR was conducted on the Applied Biosystems Step OnePlus Real-Time PCR Digital System as per manufacturer’s instructions.

### Genome-wide CRISPR screen

Genome-wide CRISPR screen for HCoV-OC43(WT) on A549 lung epithelial cells (Fig. 1A, B) was conducted previously using the Brunello library (Addgene #73179) [33]. CRISPR-screen for host factors required for HCoV-OC43/S(N29T/N514K/D535N) was conducted on 293T NXPE3/B4GALT7 DKO cells using the same Brunello library (lenticrisprV2-Blasticidin resistant), amplified according to depositor’s instructions. The resulting plasmid DNA was first PCR-amplified in four technical replicates using an equimolar pool of 8 staggered P5 forward primers (for generating sequence diversity) and a unique P7 reverse primer for each replicate (500nM per primer mix) to qualitatively analyze the library and adequate representation of gRNAs. For the PCR, 200ng plasmid DNA with Takara PrimeStar GXL DNA polymerase (#R050A) with an initial denaturation at 98°C/5 minutes followed by 28 cycles of 98°C/15s, 55°C/15s, 68°C/30s and a final extension of 68°C for 5 minutes followed by cooling at 4°C. Afterwards, lentivirus was generated on 293T cells (6*10e6 cells/150 mm dish) by transfecting 6 µg of Brunello lenticrisprV2, 6 µg of pNL-Gag-pol and 1.2µg pVSV-G using Lipofectamine 2000 (ThermoFisher #11668019) as per manufacturers’ instructions. The media was changed the next day and 24 hours later, lentivirus was harvested using 0.2-micron filter and concentrated using the Lenti X concentrator reagent (Takara #631231) as per manufacturer’s protocol. The virus pellet was resuspended in serum-free DMEM. For titration, 40,000 293T NXPE3/B4GALT7 DKO cells were transduced with serial dilutions (10-fold) of lentivirus and subject to 5 µg/ml Blasticidin selection 48 hours later. A suitable dilution to count clonal populations was used to determine the lentivirus titer. Later, 1.3*10e8 293T NXPE3/B4GALT7 DKO cells were transduced at an MOI:0.3 using polybrene (8 µg/mL) and selected using Blasticidin 48 hours later for 7-10 days. Thereafter, 8*10e6 293T NXPE3/B4GALT7cells were either left uninfected or infected with HCoV-OC43/S(N29T/N514K/D535N) at an MOI:0.1, in triplicates. 2 weeks post-infection, cellular genomic DNA was extracted using Zymo Quick-DNA midi prep Plus kit according to manufacturer’s protocol. Libraries were generated by PCR using Takara PrimeStar GXL polymerase in 100uL reactions using 10 µg of genomic DNA with an initial denaturation at 98°C/5 minutes followed by 28 cycles of 98°C/15s, 55°C/15s, 68°C/30s and a final extension of 68°C for 5 minutes followed by cooling at 4°C. Amplification was done using an equimolar pool of 8 staggered P5 forward primers and a unique P7 reverse primer for each replicate at a final concentration of 500nM per primer set. PCR products were then gel purified after 1% agarose gel electrophoresis using the Nucleospin Gel & PCR Clean up kit (Macherey-Nagel # 740609.250). All samples were pooled and sequenced using NextSeq 2000 (1×100 bp single-ended sequencing) at the Rockefeller University Genomics Resource Center. Information on all Illumina P5 and P7 primers used can be found in the Supplementary Table 2.

### Establishment of stable cell lines

#### Plasmid cloning

All plasmids are listed in the Supplementary Table 2. All knockout cell lines were generated by CRISPR-cas9 using the lenticrisprV2 plasmid (Addgene #52961) (with hygromycin/puromycin/Blasticidin/bleomycin mammalian resistance genes) that encodes both Cas9 and a U6-driven gRNA. Cloning was performed using the plasmid depositor’s protocol. Briefly, gRNAs were generated by annealing & phosphorylation of both forward and reverse oligos purchased from IDT as per depositors’ instructions using T4 PNK enzyme (NEB #M0201). The lenticrisprV2 plasmid was digested using BsmBI-v2 (NEB #R0739) and annealed oligos were cloned with T4 ligase (NEB #M0202) using manufacturer’s protocol. Ligation was performed at room temperature followed by transformation.

All plasmids for stable protein expression were generated on the pLX304 lentiviral plasmid (CCIH lentiviral plasmid with hygromycin resistance was used for ACE2 expression) which uses a CMV promoter to drive expression of the gene of interest or using the CSIH/P/B/N (Hygromycin/puromycin/blasticidin/Neomycin resistance) which uses the SFFV promoter. All genes were cloned via Gibson assembly between the BsrGI and NheI sites (for pLX304) or between BamHI and/or SfiI (for CSIH/P/B/N) using the 2X Gibson Assembly Master Mix (NEB #E2611). Gene coding sequences were purchased as gene blocks from IDT and sequences of NXPE3, B4GALT7 and IFITM2/3 were codon-optimized to make them resistant to guide RNA. Individual mutations for NXPE3 and IFITM proteins were made via PCR-directed mutagenesis followed by cloning using Gibson assembly. Assembly was performed at 50°C for 20-30 minutes followed by transformation. PCR products for cloning were amplified using Takara PrimeStar GXL polymerase with an initial denaturation at 98°C/5 minutes followed by 35 cycles of 98°C/15s, 55°C/15s, 68°C/30s (per kb) and a final extension of 68°C for 5 minutes followed by cooling at 4°C. All PCR products and/or linearized plasmids were extracted by 1% agarose gel electrophoresis followed by gel purification using the Nucleospin Gel & PCR Clean up kit (Macherey-Nagel #740609.250). All restriction enzymes were obtained from NEB and used as per manufacturer’s protocol. Information on all recombinant DNA and plasmids used can be found in the Supplementary Table 2.

#### Transformation and verification of clones

Gibson assembly or ligation mixtures were directly transformed into homemade DH10B chemically competent cells by heat shock at 42°C for 45 seconds followed by incubation on ice (5 minutes). Bacteria were revived using SOC media for 1 hour at 30°C followed by plating on LB-Ampicillin plates at 30°C, overnight. Colonies were picked the next day and grown for plasmid mini prep. Correct clones were confirmed by nanopore sequencing (Plasmidsaurus) and/or Sanger sequencing (Psomagen). Highly concentrated plasmid DNA was obtained by Midi prep using PureYield Plasmid Midiprep System (Promega #A2495) as per manufacturers’ protocol.

#### Generation of lentivirus and transduction

Lentiviruses were generated by transfecting 4*10e5 293T cells using the lentiviral plasmid of interest, pVSV-G and pNL-Gag-pol (Ratio:1200ng:1200ng:200ng) with PEI (8.8 uL per transfection mix) in serum-free DMEM. Transfection mix was incubated for 10 minutes at RT prior to transfection. Media was changed the next day, and lentiviruses were collected 24 hours after using a 0.2-micron filter and either frozen (-80°C) or immediately used for transduction. Cells were subject to selection with antibiotics 48 hours later and maintained as a polyclonal population or used to select single-cell clones by limiting dilution where required.

#### Verification of cell lines

Cell lines expressing various proteins (pLX304 system) were screened by immunoblotting for the protein/peptide tag of interest (See Supplementary Table 2 for antibodies used). CRISPR-cas9 KO cells (lenticrisprV2 system) were screened by Sanger sequencing the locus of interest (NXPE3, B4GALT7, EXT2). Knockout efficacy was computed by the Synthego ICE software, while IFITM 1/2/3 knockout cells were screened by immunoblotting for IFITM2 and IFITM3.

### Small-scale CRISPR screen

Small-scale screening for host factors that determine entry of HCoV-OC43/S(N29K) virus was conducted. Based on hits obtained in **Fig. 1A**, we focused on proteins that localize to the membrane or have unknown intracellular localization. We prepared lentiviruses (lenticrisprv2-puro) and generated polyclonal 293T cell lines for CRISPR knockout as described under establishment of stable cell lines. Two sgRNAs for each gene targeting CASD1, SLC35B2, B4GALT7, LAMTOR2, CCDC93, WRD81, SEC24C, CCDC22, KCNMA1, WDR91, PXTLP1, XYLT2, CT47A5, and PIK3C3 were cloned. Cell lines were then infected with HCoV-OC43/S(N29K) at an MOI:0.1 and infectivity was assessed by flow cytometry to assess host factors required for HCoV-OC43/S(N29K) virus. Sequences of all gRNAs used can be found in the Supplementary Table 2.

### Establishment of HCoV-OC43 and HCoV-229E reverse genetics platform (CPER)

#### Generation of plasmids for circular polymerase extension reaction (CPER)

The HCoV-OC43 and 229E reverse genetics systems were based on the previously published circular polymerase extension reaction (CPER) methodology for SARS-CoV2 [35], which was used to generate SARS-CoV-2 CPER system. Improved CPER was utilized for HCoV-229E [62]. Briefly, 3-5 kb PCR fragments (8-11) spanning the entire viral genome of OC43 ATCC VR-1588 and 229E ATCC VR-740 were cloned into a lentiviral CSIN vector between BamHI and SfiI sites using Gibson assembly (See Supplementary Table 2 for primer sequences). These PCR fragments were amplified from ATCC virus stocks. Briefly, viral RNA was extracted using the Nucleospin RNA virus kit (Macherey-Nagel) as per protocol. cDNA was then prepared using SuperScript III Reverse Transcriptase (Thermo Fisher # 18080044) which was used as a template to amplify PCR fragments using Takara PrimeStar GXL polymerase. An additional linker plasmid was cloned which contained homologies to the first and last fragment of the viral genome (See Fig. S2). Cloning was performed as described under plasmid cloning. Correct clones were confirmed by nanopore sequencing (Plasmidsaurus) and/or Sanger sequencing (Psomagen). Highly concentrated plasmid DNA was obtained by Midi prep using Promega PureYield Plasmid Midiprep System as per manufacturers’ protocol. Generation of individual mutations at OC43 spike was done via Gibson assembly and PCR-directed mutagenesis and cloning the respective plasmid. GFP reporter virus for HCoV-OC43 was generated by replacing the ns2 ORF with M protein C-terminally fused to GFP and cloning the respective plasmid. HCoV-229E GFP virus was generated by inserting GFP downstream of a TRS present immediately after the nucleocapsid ORF and cloning the respective plasmid. SARS-CoV2 GFP reporter virus was generated by replacing ORF 7a/b by the GFP coding sequence and cloning the respective plasmid. Plasmids and oligos used for the SARS-CoV-2 CPER system were obtained from [35]. Information on all plasmids and sequences of all oligos used for cloning the plasmids (CSIN primers) and for the CPER (CPER primers) can be found in the Supplementary Table 2.

#### Generation of PCR amplicons and CPER assembly

Individual PCR fragments for HCoV-OC43 and 229E CPER were generated by using CPER primers (see Supplementary Table 2). These primers introduced 40-50bp homologies between each adjacent fragment in the virus genome. Amplification was conducted using Takara PrimeStar GXL polymerase. PCR products were then gel purified after 1% agarose gel electrophoresis using the Nucleospin Gel & PCR Clean up kit (Macherey-Nagel). For the CPER reaction, 50 uL PCR reactions using the Takara PrimeStar GXL polymerase enzyme was conducted which included 10uL 5X GXL buffer, 4uL dNTP mix, 1 uL of enzyme and 100 ng of each of the fragments for the respective virus. Volume was adjusted to 50 uL using nuclease-free water. Conditions used were as follows: Initial denaturation at 98°C/5 minutes followed by 35 cycles of 98°C/15s, 55°C/15s, 68°C/25 minutes and a final extension of 68°C for 30 minutes followed by cooling at 4°C.

For HCoV-229E, an improved CPER protocol was utilized [62]. Prior to CPER, individual PCR fragments were subject to phosphorylation by T4 PNK (NEB) and re-purified using the Nucleospin Gel & PCR Clean up kit (Macherey-Nagel), followed by CPER reaction. After CPER, the mix was subject to ligation using 48 uL CPER mix with 1uL 50mM NAD^+^ (NEB #B9007) and 1ul HiFi Taq DNA ligase (NEB #M0647). The reaction was incubated at 50°C for 30 minutes followed by 60°C for 30 minutes and cooling at 4°C, ready to transfect.

#### Transfection and generation of virus stocks

For virus reconstitution, 25 uL of CPER mix was incubated with 8 uL PEI in serum-free DMEM for 10-15 minutes and directly transfected on 293T (HCoV-OC43), 293T-hAPN (HCoV-229E) or 293T-ACE2 cells (SARS-CoV2). Upon obtaining complete cytopathic effects, cell-free supernatant was collected and used to infect a new batch of the above-mentioned cells to generate crude passage 1 stock. For SARS-CoV-2, this was done on Vero E6 cells to allow better propagation of virus.

### Purification of BCoV-HE-Fc His tagged protein

BCoV-HE (19 to 377 aa; NCBI accession AAA66393) with a mutation at S40A [serine (AGC) to alanine (GCC) at 40 aa] for enzymatically inactive protein [39] was fused with IgG Fc region (IGHG1) and 6 x His-tag to its C-terminus, was cloned into the pCAGGS expression vector as described previously [29]. BCoV-HE protein was produced using the Expi293 Expression System according to the manufacturer’s protocol. In briefly, 25 µg of the above expression plasmid was transfected into 7.5 x 10E7 Expi293T cells using ExpiFectamine 293. The following day, Enhancer1 and 2 were introduced, and the cells were further cultured for an additional four days. On the day five after plasmid transfection, the culture supernatant was collected, filtered, and then incubated overnight at 4°C with Ni-NTA agarose beads (Qiagen #30210). They were collected by gravity flow into a column (BioRad # 7372522) and then washed with 20mM imidazole wash buffer (50mM NaH2PO4, 300mM NaCl, 20mM Imidazole, pH:8.0), the proteins were eluted with 250 mM imidazole. Eluted proteins were dialyzed into 1X PBS using Slide-A-Lyze Dialysis Cassettes, 10K MWCO (ThermoFisher #66380). Concentration of protein was measured using varying known amounts of BSA as standard by SDS-PAGE on a NuPAGE Bis-Tris Mini Protein Gels, 4–12%, 1.0–1.5 mm (Invitrogen) and both amount and purity was assessed by Coomassie brilliant blue staining (GelCode Blue Safe Protein Stain; ThermoFisher #24596).

### Flow cytometry assays

All flow cytometry experiments were conducted on the Attune NxT Cytometer (Acoustic focusing cytometry) and Attune CytKick Autosampler according to manufacturer’s instructions and data was acquired on the Attune Cytometric software v6.2.1.

#### Infection assays

Infections were conducted on 96-well flat-bottom plates using 2*10e4 cells/well for infection with viruses at different MOIs in DMEM media supplemented with FCS and Gentamicin. At a suitable time point (see figure legends for each experiment), Cells were washed in 1X PBS, Trypsinized and fixed in 4% PFA for 15 minutes. Cells were washed and permeabilized with 0.5 % Triton X-100 for five minutes followed by washing. Cells were either directly processed for flow cytometry when using GFP reporter viruses or stained with antibodies against HCoV-OC43/ HCoV-NL63 nucleocapsid for 1 hr at RT. Cells were washed and incubated with secondary Ab Alexa fluor 488 goat anti-mouse IgG (H+L) (Invitrogen # A-11001) for 1 hr at RT. All cells were washed and resuspended in 0.1% Poloxamer 188 Non-ionic Surfactant (Gibco # 24040032) in 1X PBS for flow cytometry analysis. All antibodies were used at 1:1000 dilution in 10% FCS in 1X PBS. 1-2 washes between each incubation step were conducted using 1X PBS by centrifugation at 500g/5 minutes. Details on experiments that involved either antibody or heparin treatment are described in the respective figure legends. Information on all antibodies used can be found in the Supplementary Table 2.

#### Detection of heparan sulfate, O-acetylated sialic acids and lectin binding

For detection of heparan sulfate and/or sialic acids cells were fixed in 4% PFA for 10-15 minutes after detachment with 1X PBS-500mM EDTA and washed. For heparan sulfate staining, cells were incubated with mouse anti-HS (Amsbio clone F58-10E4 # 370255-S) for 1 hr at RT. Cells were washed and incubated with secondary Ab Alexa fluor 488 goat anti-mouse IgG (H+L) (Invitrogen) for 1 hr at RT. For O-acetylated sialic acid detection, enzymatically inactive BCoV-HE specific for acetylated sialic acid was used. The protein was incubated at 1µg/mL on A549-based cell lines at 4°C for 1 hr followed by His-tag staining Mouse anti-His Tag Alexa Fluor 488-conjugated Antibody (R&D Systems #IC050G) at 4°C for 1 hr. Lectin binding assay was performed by using 1:1000 dilution of biotinylated *Maackia amurensis* lectin II (Mal II) (Vector Labs, B-1265) incubated at 4°C for 1 hr, followed by staining with Streptavidin Alexa fluor 488 conjugate (Invitrogen # S32354). Similarly, mouse splenocytes were stained after obtaining the cells by homogenizing spleens and resuspending filtered cells in RPMI media (Gibco # 11875119). All cells were washed and resuspended in 0.1% Pluronic F-68-1X PBS for flow cytometry analysis. All antibodies were used at 1:1000 dilution in 10% FCS in 1X PBS. 1-2 washes between each incubation step were conducted using 1X PBS by centrifugation at 500g/5 minutes.

### Microscopy

All microscopic imaging was performed on the DeltaVision OMX SR- Super Resolution Microscope (Cytiva imsol- image solutions) using the AcquireSR software with a 60X widefield oil immersion lens (Olympus). Exposure was adjusted to 100 ms and laser power was kept constant for each experiment, varying between 0.2-10% depending on target brightness. Cytiva softWoRx tools were used to align and deconvolve raw images to obtain high resolution 3D image files.

#### Virion binding assays

293T-based cell lines were seeded at 2*104 cells on 8-well chamber plates (Labtek #12-565-338 or Ibidi #80807) which were coated with Poly-D-lysine Hydrobromide (Advanced Biomatrix #5175) 1 hour prior to seeding. The next day, cells were infected with HCoV-OC43 ýns2 OC43 M-eGFP for 1.5 hr at 4°C followed by gentle 1X PBS wash and 4% PFA fixation (15 minutes). Cells were then counterstained for nuclei using DAPI and Hoechst Nucleic Acid Stains (# Invitrogen D1306 at 1 µg/ml) and for 10 minutes at RT followed by washing and imaging.

#### Immunostaining

293T-based cell lines were seeded at 2*104 cells on 8-well chamber plates (Labtek #12-565-338 or Ibidi #80807) which were coated with Poly-D-lysine Hydrobromide 1 hour prior to seeding. The next day cells were fixed in 4% PFA (15 minutes) and permeabilized in 0.5 % Triton-X 100 (5 minutes). Cells were then incubated with antibodies against various proteins including IFITM2/3 (rabbit polyclonal anti-IFITM3, Proteintech #11714-1-AP), IFITM1 (mouse monoclonal anti-IFITM1, Proteintech # 60074-1-Ig) and HA (mouse monoclonal anti-HA.11; Biolegend #901501) at RT for 1 hour. Cells were then incubated with secondary antibodies secondary Ab Alexa fluor 488 goat anti-mouse/rabbit IgG (H+L) (Invitrogen # A-11001/ # A-11008) and DAPI and Hoechst Nucleic Acid Stains (# Invitrogen D1306 at 1 µg/ml) for 1 hour at RT followed by washing and imaging. Information on all antibodies used can be found In the Supplementary Table 2.

### Immunoblot

For immunoblotting, samples were prepared in 1X NuPAGE LDS Sample buffer (Invitrogen NP0008) supplemented with 2.5 % ß-mercaptoethanol. Samples were sonicated and heated at 70°C for 10 minutes. Samples were directly loaded on NuPAGE 4-12% Bis-Tris gels (Invitrogen NP0329) or 12% Bis-Tris gels (Invitrogen NP0342). SDS-PAGE was conducted using XCell *SureLock* Mini-Cell (Invitrogen EI0001) followed by wet transfer on 0.45um nitrocellulose membrane (Cytiva Amersham Protram 0.45um NC #10600002) using XCell II Blot Module (Invitrogen EI0002) as per manufacturer’s conditions. Following transfer, membranes were blocked with Intercept PBS blocking buffer (Licorbio #927-70001) for 45 minutes at RT, followed by antibody incubations for one hour at RT or at 4°C/overnight. Between incubations, membranes were washed with 1XPBS-0.1% Tween-20. All primary antibodies were diluted 1:1000 in Intercept PBS blocking buffer and 1:2500 for secondary antibodies. Primary antibodies included antibodies against IFITM3 (detects both IFITM2/3), IFITM1, Syndecan-1, Syndecan-2, hAPN (CD-13), HA tag epitope, tubulin, ß-actin and a in-house 5B4 antibody against hACE2. Near-infrared fluorescent Secondary antibodies were obtained from Licorbio (IRDye). Imaging was performed on Licor Odyssey M Imaging system as per manufacturer’s instructions using the Licor 2.2 Acquisition software. All antibodies are listed in the Supplementary Table 2.

### Focus formation assay

Focus formation assays were conducted on 24-well plates. 4*10e5 293T-based cell lines seeded using Poly-D-lysine Hydrobromide were infected with 200 ul of 10-fold virus serial dilutions for 1 hour at 34°C at 5% CO_2_ with gentle shaking every 15 minutes. Media was replaced with 1% carboxy-methyl cellulose (CMC) overlay comprising DMEM media supplemented with 2% FCS and gentamicin with 1% CMC and cells were incubated for 4-5 dpi. At 4-5 dpi cells were either fixed with 4% PFA and immunostained for viral nucleocapsid proteins as described before or directly visualized for GFP expression. Cells were imaged at 4X magnification on the Invitrogen EVOS M7000 Imaging System and both individual and tiled images of each well were collected. Foci were manually counted at a suitable dilution range to determine viral titers. All antibodies are listed in the Supplementary Table 2.

### Live-cell NanoLuc luciferase assays

Live-cell NanoLuc assays to measure virus fusion and primary translation were conducted on 2*10e4 cells/well seeded the prior day on black, flat bottom 96-well plates (CoStar). The cells were infected with either the HCoV-OC43_M-HiBiT (fusion) or the HCoV-OC43_nsp3-Nluc viruses in the presence of 1:100 dilution of Endurazine reagent (Promega) with or without 50ug/ml of cycloheximide or DMSO solvent control. Luminescence values were measured at various time points on the GloMax Navigator system (Promega)

### Quantification and statistical analysis

For CRISPR-screen analysis, raw fastq files obtained after high-throughput Illumina sequences were analyzed on the Galaxy open-source project. Briefly, demultiplexed and adaptor-trimmed reads (Cutadapt) based on barcoded P7 primers were subject to MAGeCK count (default settings) to assess the number of gRNAs sequenced against the Brunello library gRNA sequences. Plasmid DNA quality of the Brunello library and gRNA distribution was first computed, giving a GiniIndex score <0.1 indicative of a highly equal distribution of gRNAs. Accordingly, the genome-wide CRISPR-screen experiment was performed. After MAGeCK count, MAGeCK test (default settings) was performed to obtain both normalized read counts and robust rank aggregation (RRA) scores to identify top host factors required for HCoV-OC43/S(N29T/N514K/D535N) infection based on fold change enrichment and p-values as listed in **Supplementary table 1**. Publicly available adapter-trimmed and QC-adjusted RNA-seq data for cell lines including 293T (SRR8950022/23), HuH7 (SRR23268608/9), HuH1 (SRR5381417), A549 (ERR12551547) and VeroE6 (SRR12164495) were analyzed on the Galaxy open-source project. Briefly, RNA-seq data was aligned against the hg38 human genome using RNA-STAR. Reads were counted for each gene using exon annotations with FeatureCounts. Counts were analyzed for differential gene expression to obtain library and gene length-normalized reads per kilobase per millions (RPKM) values using DeSeq2. Heat-map visualization of RPKM read counts was performed on Prism 11. All flow cytometry data was analyzed using FloJo 10. Confocal microscopy 3D images were visualized on Imaris for scale bar calibration. Focus formation assay data was analyzed on Fiji to measure plaque size by free-hand shaping tools. All tabular data obtained for all experiments were analyzed on Prism 11 to generate graphs and perform statistical analyses. Immunoblot results were analyzed on ImageStudio.

## Resource availability

### Lead contact

Further information and requests for resources should be directed to and will be fulfilled by the lead contacts

### Materials availability

New reagents described herein are available on request.

### Data and code availability

CRISPR screening data is available in the supplementary table

## Acknowledgments

This work was supported by a grant from the Stavros Niarchos Foundation, The Rockefeller University and by the Howard Hughes Medical Institute. We thank the Rockefeller University flow cytometry core, Comparative Biosciences Center (CBC), and Genomic Resource Center for their contributions. This article is subject to HHMI’s Open Access to Publications policy. HHMI lab heads have previously granted a nonexclusive CC BY 4.0 license to the public and a sublicensable license to HHMI in their research articles. Pursuant to those licenses, the author-accepted manuscript of this article can be made freely available under a CC BY 4.0 license immediately upon publication.

## Author contributions

PDB, TH, ML and MI conceived the project. ML and MI performed the experiments with the help of MT, MC, DP and YW. NAS and SP generated and advised on the use of HCoV-OC43_NV_. PDB and TH supervised the project. ML and MI wrote the manuscript. PDB, TH and SP revised and edited the manuscript.

---

**Fig. S1:**
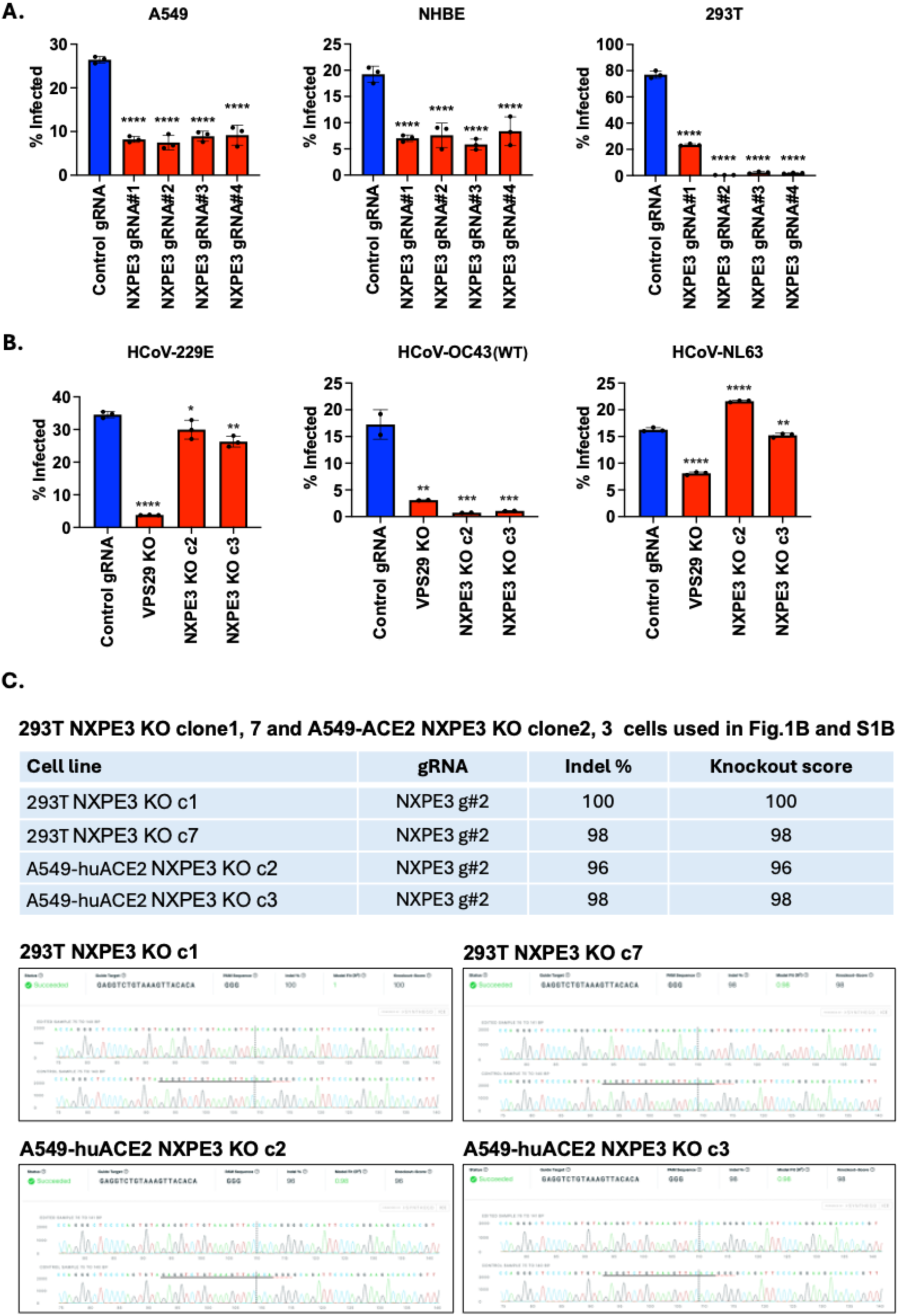
NXPE3 Knockout efficacy and effects on HCoV-OC43 infection. (A) A549, NHBE and 293T cells transduced with a control or NXPE3-targeting guide RNAs were infected with HCoV-OC43(WT) for 24 h. Infection was assessed by flow cytometry using antibodies against viral nucleocapsid proteins. Error bars indicate SEM; each dot represents a replicate. P-values are determined by one-way ANOVA (****, p<0.0001) (B) Control and single-cell NXPE3/VPS29 knockout A549-ACE2 cells were infected with various coronaviruses for 24 h. Infection was assessed by flow cytometry using antibodies against viral nucleocapsid proteins. Error bars indicate SEM; each dot represents a replicate. P-values are determined by one-way ANOVA (****, p<0.0001; ***, p<0.001; **, p<0.005; *, p<0.05) (C) Gene knockout efficacy determined by Sanger sequencing and Synthego ICE software for cell lines used in Fig 1 and S1 are shown.

**Fig. S2:**
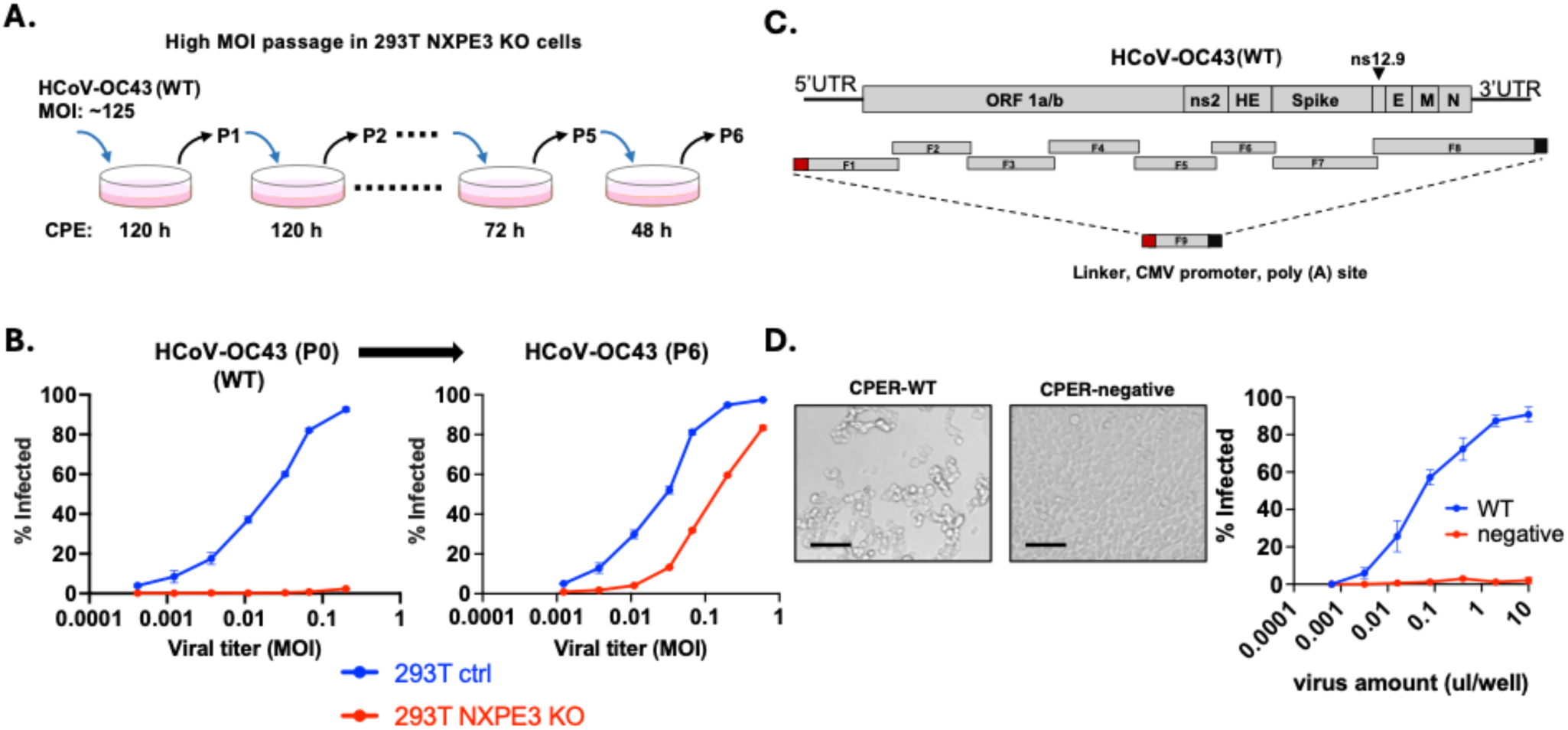
Spike N29T mutation enables NXPE3-independent HCoV-OC43 infection. (A) Experimental schematic of high MOI passaging of HCoV-OC43(WT) on 293T NXPE3 KO cells to obtain NXPE3-independent HCoV-OC43, isolated by six passages as shown in (B) where infectivity of titrated supernatants was measured by flow cytometry. Error bars represent SEM (n=3 per MOI). (C) Schematic representation of the HCoV-OC43 CPER reverse genetics system as was done for SARS-CoV-2. (D) Left: Cytopathic effects of HCoV-OC43-infected 293T cells obtained in CPER-WT samples. CPER-negative represents CPER reaction performed with 1-2 missing fragments. (D) Right: HCoV-OC43 infection efficacy from CPER-WT and CPER-negative samples on 293T cells using various volumes of cell-free supernatant. Infection was assessed by flow cytometry using antibodies against viral nucleocapsid proteins. Error bars represent SEM (n=2 per MOI).

**Fig. S3:**
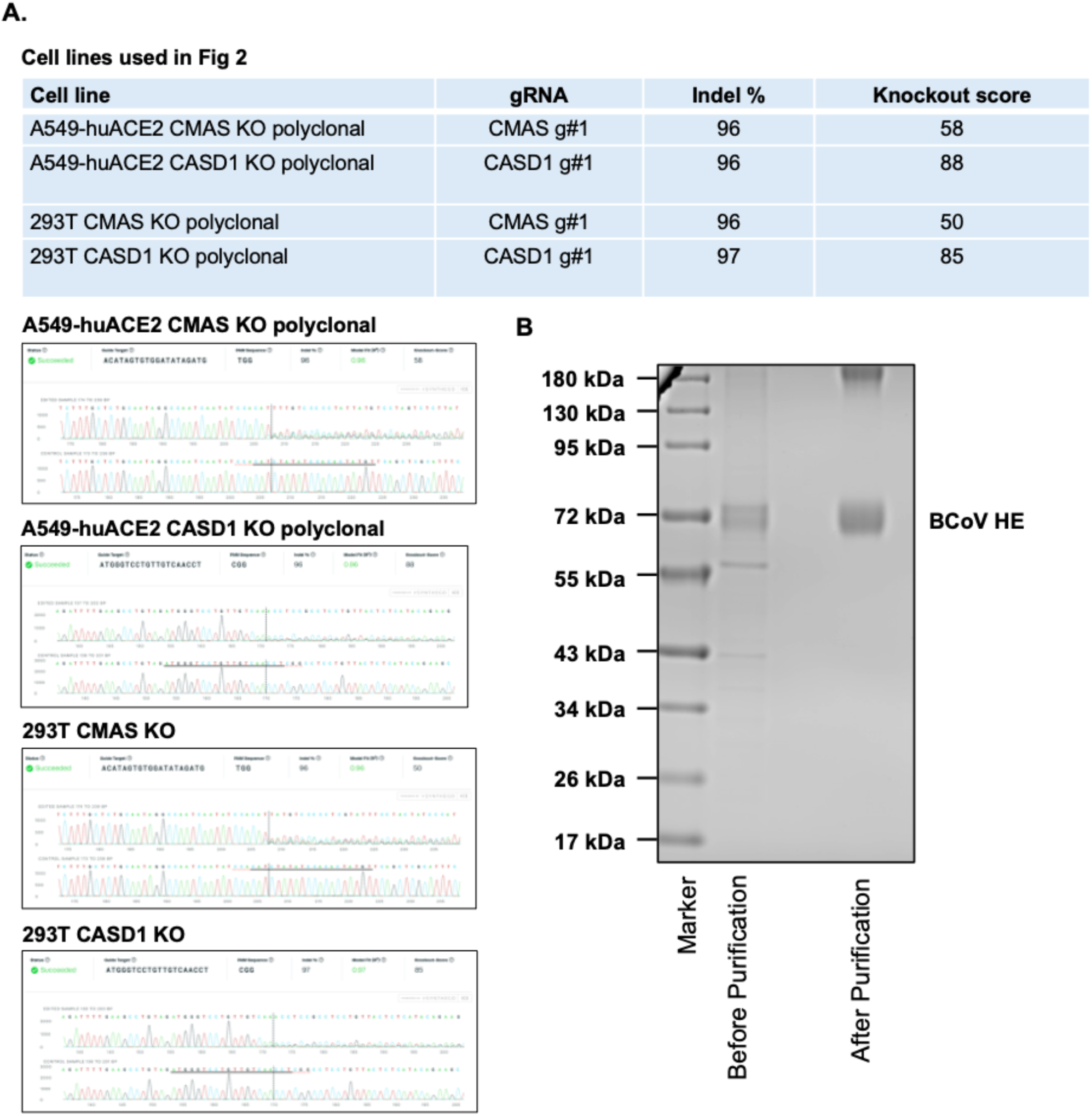
Detection of BCoV-HE protein reagent and CRISPR-cas9 knockout for synthesis of 9-O-Ac-Sia in 293T and A549-ACE2 cells. (A) CRISPR-cas9 knockout efficacy for CMAS and CASD1 in 293T and A549-huACE2 cells as determined by Sanger sequencing and Synthego ICE software. (B) Coomassie brilliant blue staining for purified BCoV-HE protein fused to an Fc domain and a C-terminal His tag for Ni-NTA purification

**Fig. S4:**
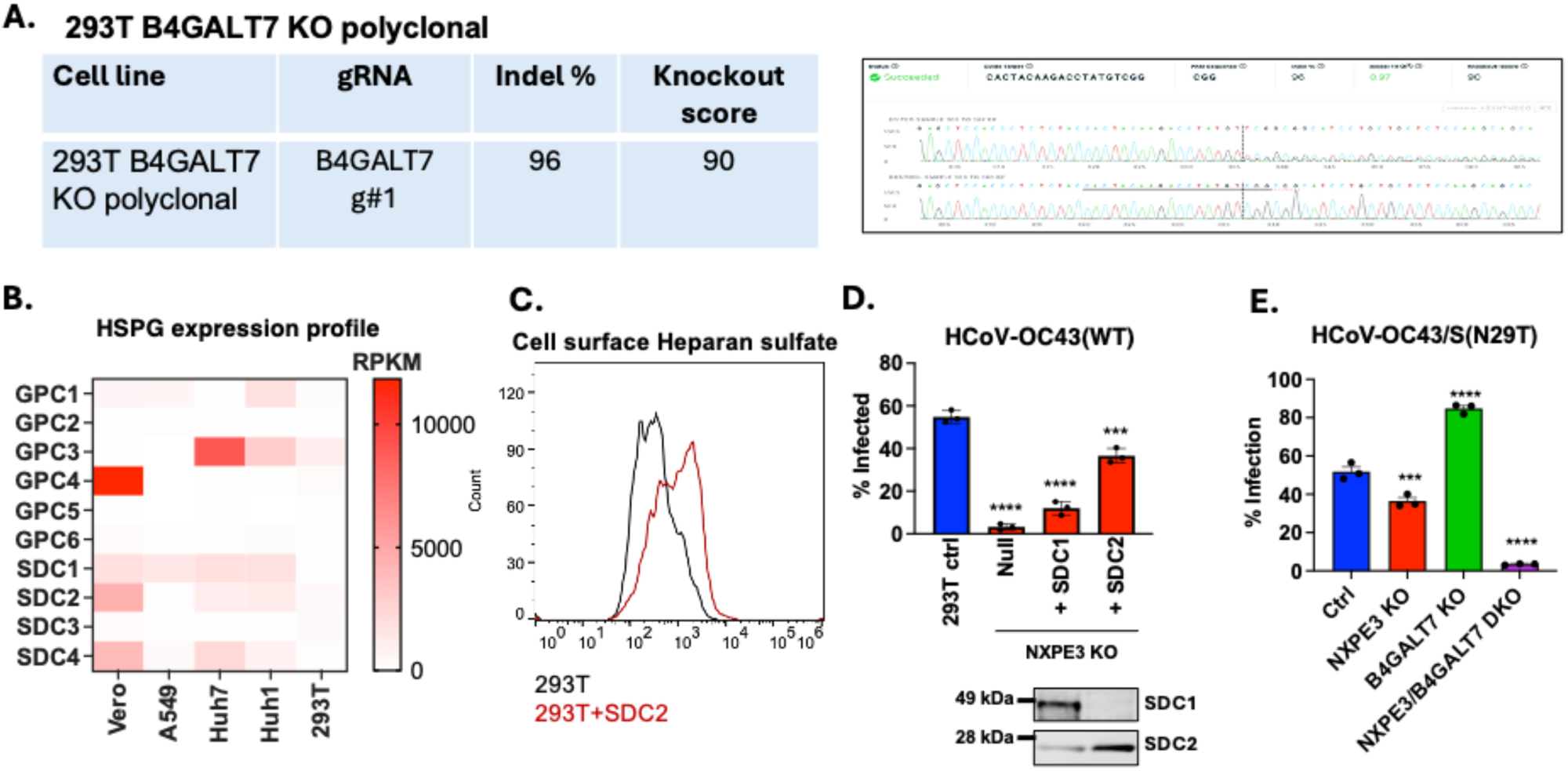
HCoV-OC43/S(N29K) infection depends on heparan sulfate. (A) Gene knockout efficacy determined by Sanger sequencing and Synthego ICE software for cell lines used in Fig 4 are shown. (B) RPKM values reflecting gene expression (high: red, low: white) for Syndecans and Glypican HSPGs across various cell lines shown as a heatmap. RNA-seq data was obtained from public datasets, as described in methods. (C) HEK293T cells transfected with SDC2 or empty vector were fixed and stained for cell-surface heparan sulfate. HS levels are depicted (x-axis) as a histogram for a representative replicate. (D) Cell lines (293T-based) were infected with HCoV-OC43(WT) for 24 h at an MOI:0.1. Bottom panels show immunoblot protein levels of Syndecan-1 and 2. Infectivity was assessed by HCoV-OC43 nucleocapsid staining and flow cytometry (n=3, Mean values with SEM are shown). P-values are determined by one-way ANOVA (****, p<0.0001; ***, p<0.001) (E) Cell lines (293T-based) were infected with HCoV-OC43/S(N29T) for 24 hours (MOI:0.2, n=3, mean values with SEM are shown) and infectivity was assessed by HCoV-OC43 nucleocapsid staining and flow cytometry. P-values are determined by one-way ANOVA (****, p<0.0001; ***, p<0.001).

**Fig. S5:**
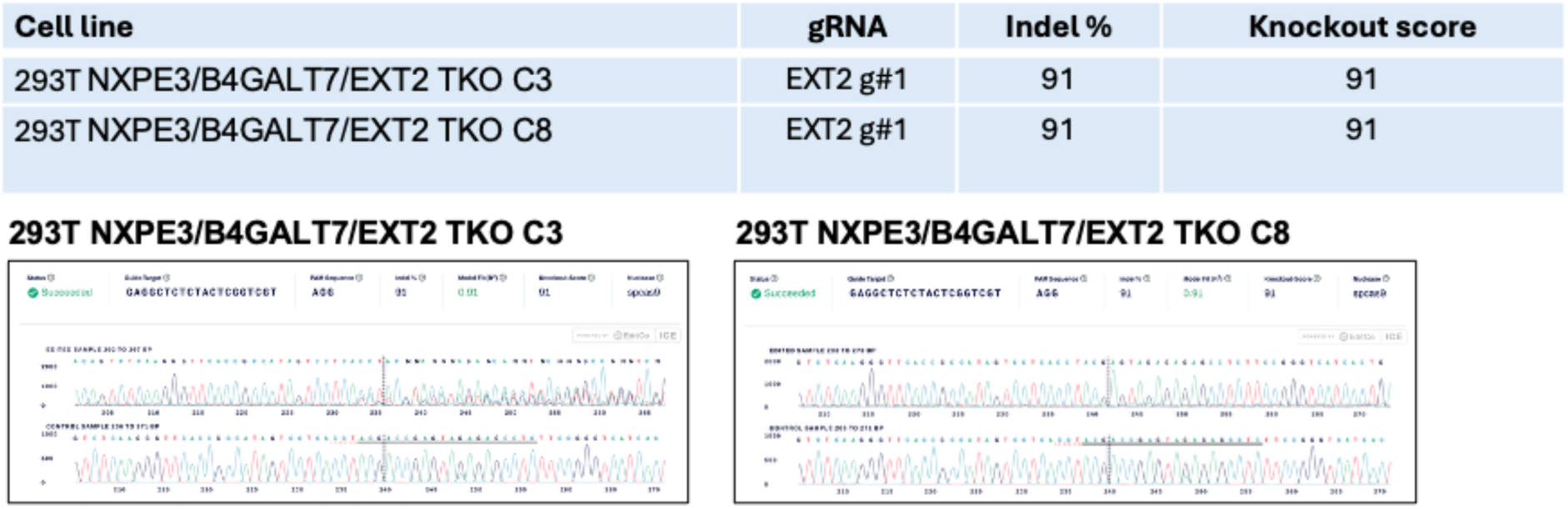
CRISPR-cas9 knockout of EXT2 in 293T NXPE3/B4GALT7 DKO cells. CRISPR-cas9 knockout efficacy of EXT2 as determined by Sanger sequencing and Synthego ICE software analysis.

**Fig. S6:**
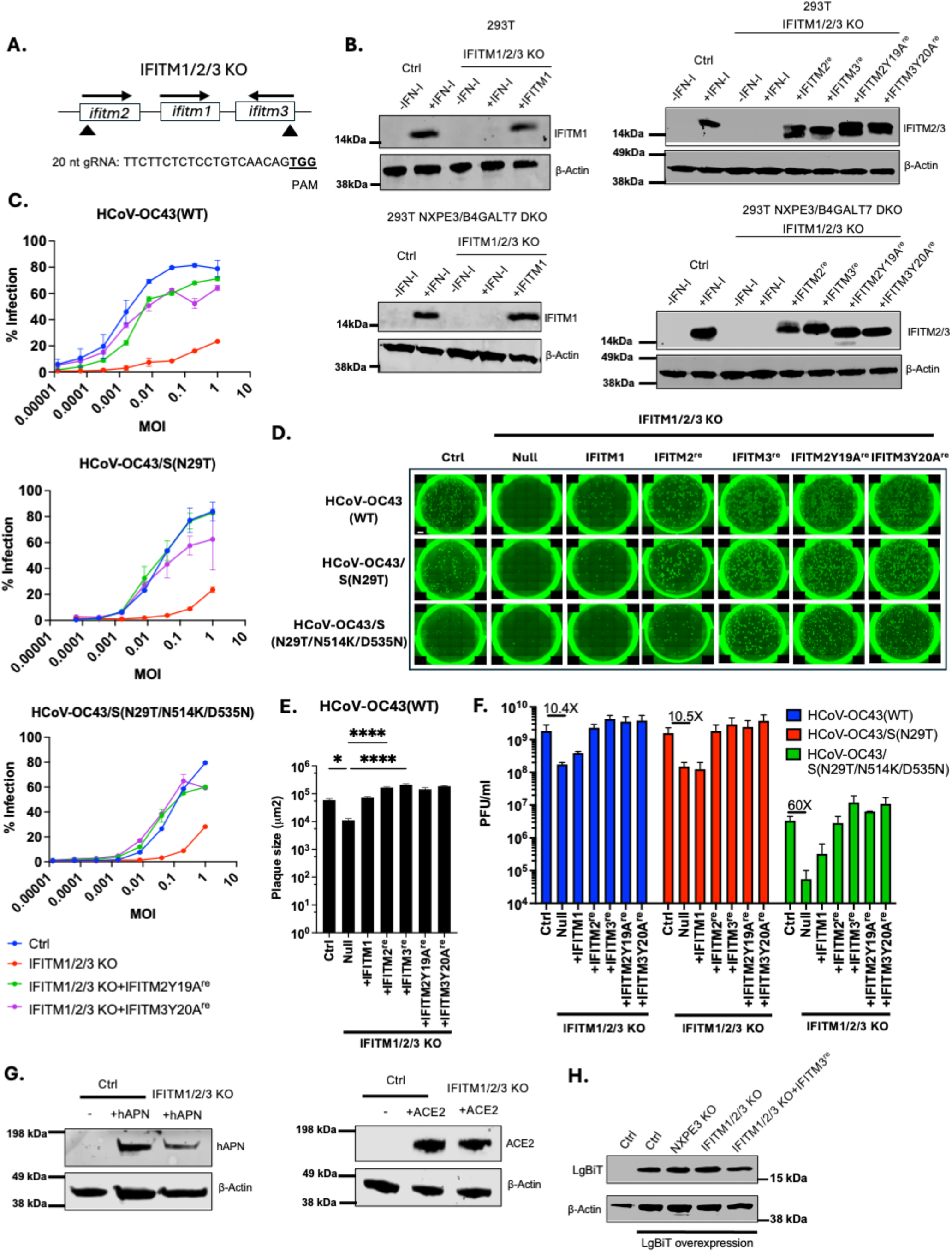
IFITM proteins are required for HCoV-OC43 infection and spike-dependent fusion, irrespective of the upstream attachment factor. IFITM1/2/3 knockout 293T cell lines were generated by using a gRNA targeting both IFITM2 and 3 (A). (B) Immunoblots showing IFITM1/2/3 knockout efficacy and IFITM expression for both 293T (upper panels) and NXPE3/B4GALT7 DKO 293T cells (lower panels) with ß-actin as loading control. The blots show IFITM (left) or IFITM2/3 (right) expression. (C) HCoV-OC43 variants were used to infect 293T-based cell lines at various MOIs for 24 h (n=2 per MOI, bars represent SEM). (D) Focus formation assays for HCoV-OC43 infection (5 dpi) on various 293T cells using 1% CMC overlay media. Representative dilutions are shown (n=2 per dilution/per virus). Foci were detected by immunostaining against N protein. Plaque sizes for 10 plaques for each infected cell line were computed in um^2^ area units for HCoV-OC43(WT) (E). SEM is shown and statistical significance was determined by one-way Anova (****, p<0.0001; *, p<0.05). (F) Viral titers (PFU/ml) obtained from focus formation assays as shown in (D) (n=2 for each virus per cell line). Error bars represent SEM. Fold change relative to control is displayed. (G) Immunoblot for hAPN and ACE2 expression for cell lines used in Fig. 6D. Immunoblot for LgBiT expression for cell lines used in Fig. 6F. Representative immunoblots are shown.

**Fig. S7:**
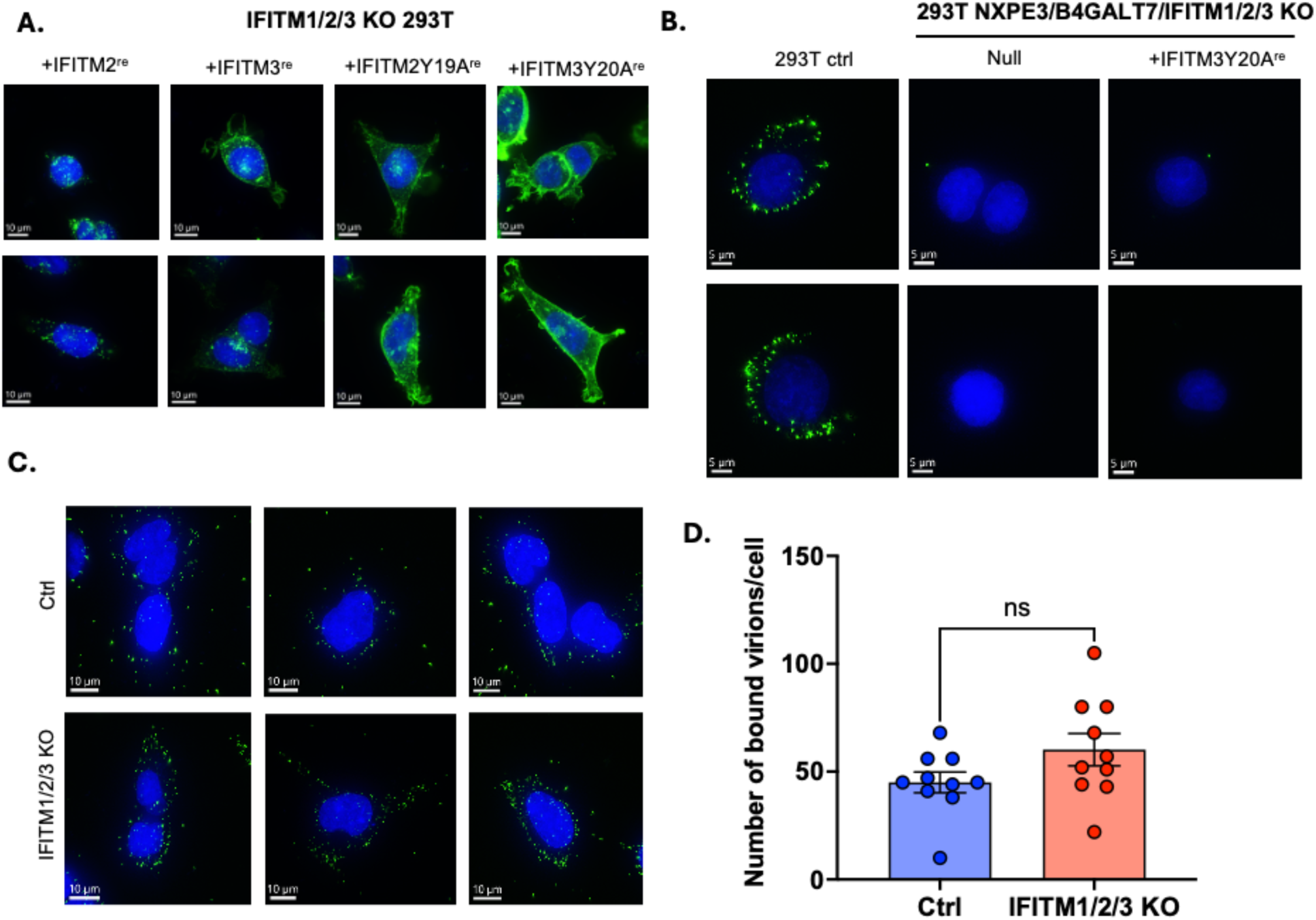
IFITM proteins are not required for HCoV-OC43 attachment. (A) Confocal images for IFITM2^re^/3^re^/2Y19A^re^/3Y20A^re^ expression (Green) on 293T IFITM1/2/3 KO cells. Hoechst (blue) used as counterstain for nuclei. (B) and (C) Virion binding assays were conducted at an MOI:20 on various 293T cells by incubating at 4°C for 1.5 hours, followed by 4% PFA fixation and microscopy. Representative images are shown. Hoechst staining (blue) used as nuclear counterstain and virions are shown as green dots. (B) No virion binding detected on 293T NXPE3/B4GALT7/IFITM1/2/3 KO cells +/-IFITM3Y20A^re^. (D) Number of bound virions/cell (10 cells/condition) for data shown in (C). Error bars represent SEM. Welch’s t-test is performed (ns: not significant).

**Fig. S8:**
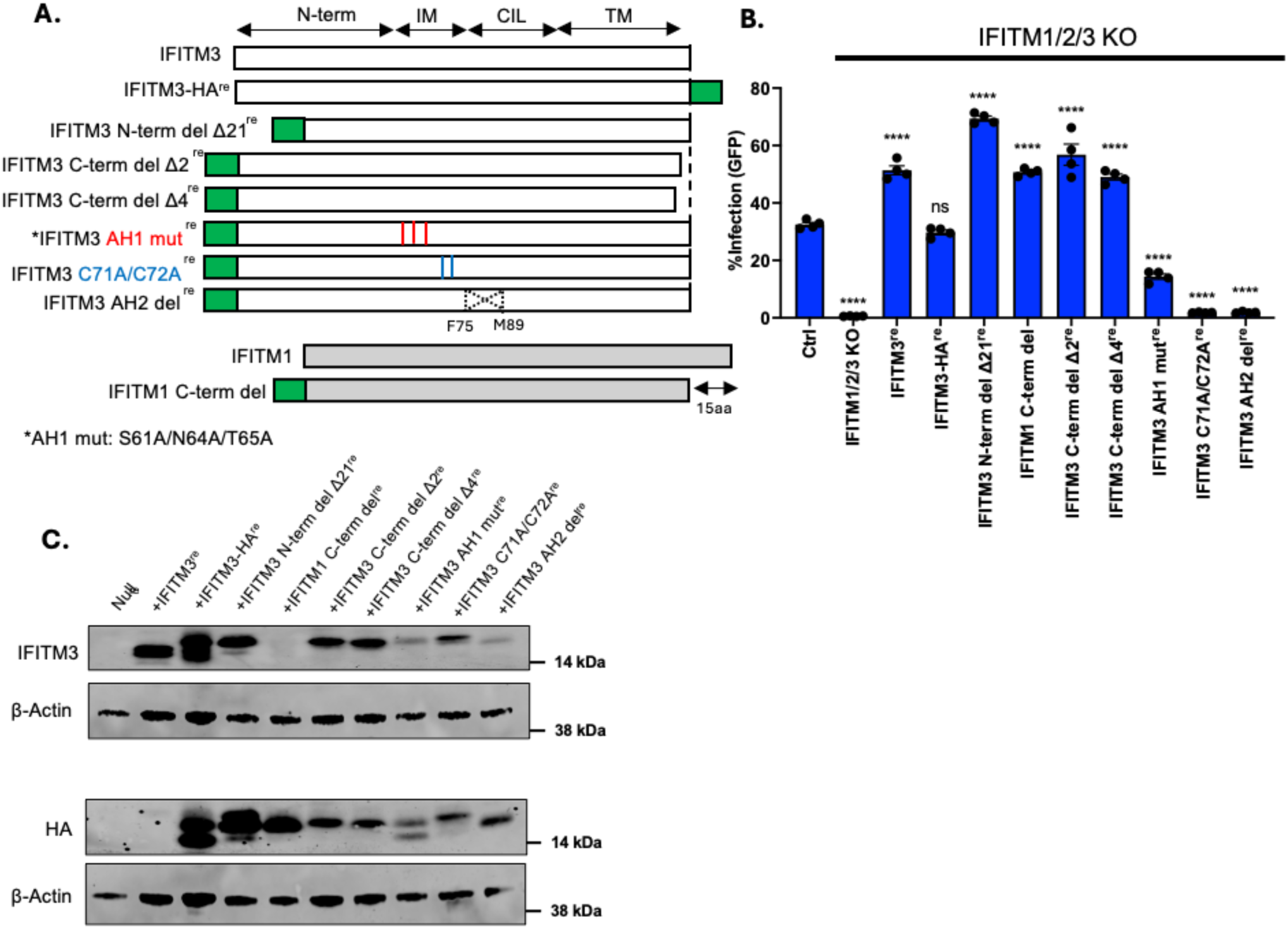
Functional domains of IFITM3 required for HCoV-OC43 infection. (A, B) Cell lines (expressing IFITM3^re^ mutants as shown in schematic in (A)) were infected with HCoV-OC43-dns2-M-GFP virus for 24 hours followed by flow cytometry to measure infectivity by GFP +ve infected cells (B). n=3 (error bars represent SEM). P-values were computed using one-way Anova (****, p<0.0001; ***, p<0.001; **, p<0.005; *, p<0.05; ns: not significant). Schematic of IFITM3 in (A) portrays the N-terminal domain, The intramembrane domain (IM) containing the amphipathic helix 1 and palmitoylation sites; The conserved intracellular loop (CIL) which contains part of the amphipathic helix 2 and the transmembrane domain (TM). (C) Immunoblot for IFITM3 and HA epitope for cell lines used in (B). ß-actin was used as loading control. Representative immunoblot is shown.

**Supplementary Table 1:** List of enriched genes obtained from MaGecK analysis for genome-wide CRISPR described in Fig. 1A and Fig. 5C. Top 10 genes are highlighted in bold. Columns C, E and F show computed p-values, gene rank and fold-change (log; comparing selection with vs without infection with HCoV-OC43 viruses).

**Supplementary Table 2:** Listing of key resources including antibodies, plasmids, primers, cell lines and viruses.

## References

1. McIntosh, K., W.B. Becker, and R.M. Chanock, Growth in suckling-mouse brain of “IBV-like” viruses from patients with upper respiratory tract disease. Proc Natl Acad Sci U S A, 1967. 58(6): p. 2268–73.

2. Vijgen, L., et al., Complete genomic sequence of human coronavirus OC43: molecular clock analysis suggests a relatively recent zoonotic coronavirus transmission event. J Virol, 2005. 79(3): p. 1595–604.

3. Butler, N., et al., Murine encephalitis caused by HCoV-OC43, a human coronavirus with broad species specificity, is partly immune-mediated. Virology, 2006. 347(2): p. 410–21.

4. Drosten, C., et al., Identification of a novel coronavirus in patients with severe acute respiratory syndrome. N Engl J Med, 2003. 348(20): p. 1967–76.

5. Zaki, A.M., et al., Isolation of a novel coronavirus from a man with pneumonia in Saudi Arabia. N Engl J Med, 2012. 367(19): p. 1814–20.

6. Chan, J.F., et al., Genomic characterization of the 2019 novel human-pathogenic coronavirus isolated from a patient with atypical pneumonia after visiting Wuhan. Emerg Microbes Infect, 2020. 9(1): p. 221–236.

7. Lu, R., et al., Genomic characterisation and epidemiology of 2019 novel coronavirus: implications for virus origins and receptor binding. Lancet, 2020. 395(10224): p. 565–574.

8. Li, F., Structure, Function, and Evolution of Coronavirus Spike Proteins. Annu Rev Virol, 2016. 3(1): p. 237–261.

9. Belouzard, S., et al., Mechanisms of coronavirus cell entry mediated by the viral spike protein. Viruses, 2012. 4(6): p. 1011–33.

10. Millet, J.K., J.A. Jaimes, and G.R. Whittaker, Molecular diversity of coronavirus host cell entry receptors. FEMS Microbiol Rev, 2021. 45(3).

11. Matsuyama, S., et al., Efficient activation of the severe acute respiratory syndrome coronavirus spike protein by the transmembrane protease TMPRSS2. J Virol, 2010. 84(24): p. 12658–64.

12. Glowacka, I., et al., Evidence that TMPRSS2 activates the severe acute respiratory syndrome coronavirus spike protein for membrane fusion and reduces viral control by the humoral immune response. J Virol, 2011. 85(9): p. 4122–34.

13. Bertram, S., et al., TMPRSS2 activates the human coronavirus 229E for cathepsin-independent host cell entry and is expressed in viral target cells in the respiratory epithelium. J Virol, 2013. 87(11): p. 6150–60.

14. Shirato, K., M. Kawase, and S. Matsuyama, Wild-type human coronaviruses prefer cell-surface TMPRSS2 to endosomal cathepsins for cell entry. Virology, 2018. 517: p. 9–15.

15. Gierer, S., et al., The spike protein of the emerging betacoronavirus EMC uses a novel coronavirus receptor for entry, can be activated by TMPRSS2, and is targeted by neutralizing antibodies. J Virol, 2013. 87(10): p. 5502–11.

16. Simmons, G., et al., Inhibitors of cathepsin L prevent severe acute respiratory syndrome coronavirus entry. Proc Natl Acad Sci U S A, 2005. 102(33): p. 11876–81.

17. Kawase, M., et al., Protease-mediated entry via the endosome of human coronavirus 229E. J Virol, 2009. 83(2): p. 712–21.

18. Li, W., et al., Angiotensin-converting enzyme 2 is a functional receptor for the SARS coronavirus. Nature, 2003. 426(6965): p. 450–4.

19. Hofmann, H., et al., Human coronavirus NL63 employs the severe acute respiratory syndrome coronavirus receptor for cellular entry. Proc Natl Acad Sci U S A, 2005. 102(22): p. 7988–93.

20. Hoffmann, M., et al., SARS-CoV-2 Cell Entry Depends on ACE2 and TMPRSS2 and Is Blocked by a Clinically Proven Protease Inhibitor. Cell, 2020. 181(2): p. 271–280.e8.

21. Raj, V.S., et al., Dipeptidyl peptidase 4 is a functional receptor for the emerging human coronavirus-EMC. Nature, 2013. 495(7440): p. 251–4.

22. Yeager, C.L., et al., Human aminopeptidase N is a receptor for human coronavirus 229E. Nature, 1992. 357(6377): p. 420–2.

23. Saunders, N., et al., TMPRSS2 is a functional receptor for human coronavirus HKU1. Nature, 2023. 624(7990): p. 207–214.

24. Vlasak, R., et al., Human and bovine coronaviruses recognize sialic acid-containing receptors similar to those of influenza C viruses. Proc Natl Acad Sci U S A, 1988. 85(12): p. 4526–9.

25. Huang, X., et al., Human Coronavirus HKU1 Spike Protein Uses O-Acetylated Sialic Acid as an Attachment Receptor Determinant and Employs Hemagglutinin-Esterase Protein as a Receptor-Destroying Enzyme. J Virol, 2015. 89(14): p. 7202–13.

26. Hulswit, R.J.G., et al., Human coronaviruses OC43 and HKU1 bind to 9-O-acetylated sialic acids via a conserved receptor-binding site in spike protein domain A. Proc Natl Acad Sci U S A, 2019. 116(7): p. 2681–2690.

27. Owczarek, K., et al., Early events during human coronavirus OC43 entry to the cell. Sci Rep, 2018. 8(1): p. 7124.

28. Tortorici, M.A., et al., Structural basis for human coronavirus attachment to sialic acid receptors. Nat Struct Mol Biol, 2019. 26(6): p. 481–489.

29. Zhang, F., et al., Pan-sarbecovirus prophylaxis with human anti-ACE2 monoclonal antibodies. Nat Microbiol, 2023. 8(6): p. 1051–1063.

30. Blachier, S., et al., Targeting ACE2 with a camelid antibody inhibits SARS-CoV-2 binding and has protective effects in vivo. Nat Commun, 2025. 16(1): p. 10268.

31. Du, Y., et al., A broadly neutralizing humanized ACE2-targeting antibody against SARS-CoV-2 variants. Nat Commun, 2021. 12(1): p. 5000.

32. Li, M., et al., Bispecific antibodies provide broad neutralization of emerging beta-coronaviruses by targeting ACE2 and viral spikes. Emerg Microbes Infect, 2024. 13(1): p. 2404166.

33. Poston, D., et al., VPS29 Exerts Opposing Effects on Endocytic Viral Entry. mBio, 2022. 13(2): p. e0300221.

34. Anantharaman, V. and L. Aravind, Novel eukaryotic enzymes modifying cell-surface biopolymers. Biol Direct, 2010. 5: p. 1.

35. Torii, S., et al., Establishment of a reverse genetics system for SARS-CoV-2 using circular polymerase extension reaction. Cell Rep, 2021. 35(3): p. 109014.

36. Jeong, J.H., et al., Molecular analysis of S gene of spike glycoprotein of winter dysentery bovine coronavirus circulated in Korea during 2002-2003. Virus Res, 2005. 108(1-2): p. 207–12.

37. Baumann, A.M., et al., 9-O-Acetylation of sialic acids is catalysed by CASD1 via a covalent acetyl-enzyme intermediate. Nat Commun, 2015. 6: p. 7673.

38. Roseman, S., Enzymatic synthesis of cytidine 5’-mono-phospho-sialic acids. Proc Natl Acad Sci U S A, 1962. 48(3): p. 437–41.

39. Barnard, K.N., et al., Expression of 9-O- and 7,9-O-Acetyl Modified Sialic Acid in Cells and Their Effects on Influenza Viruses. mBio, 2019. 10(6).

40. Wang, W.C. and R.D. Cummings, The immobilized leukoagglutinin from the seeds of Maackia amurensis binds with high affinity to complex-type Asn-linked oligosaccharides containing terminal sialic acid-linked alpha-2,3 to penultimate galactose residues. J Biol Chem, 1988. 263(10): p. 4576–85.

41. Almeida, R., et al., Cloning and expression of a proteoglycan UDP-galactose:beta-xylose beta1,4-galactosyltransferase I. A seventh member of the human beta4-galactosyltransferase gene family. J Biol Chem, 1999. 274(37): p. 26165–71.

42. Cagno, V., et al., Heparan Sulfate Proteoglycans and Viral Attachment: True Receptors or Adaptation Bias? Viruses, 2019. 11(7).

43. Ramos-Martínez, I.E., et al., Heparan Sulfate and Sialic Acid in Viral Attachment: Two Sides of the Same Coin? Int J Mol Sci, 2022. 23(17).

44. Esko, J.D., et al., Inhibition of chondroitin and heparan sulfate biosynthesis in Chinese hamster ovary cell mutants defective in galactosyltransferase I. J Biol Chem, 1987. 262(25): p. 12189–95.

45. Schneider, W.M., et al., Genome-Scale Identification of SARS-CoV-2 and Pan-coronavirus Host Factor Networks. Cell, 2021. 184(1): p. 120–132.e14.

46. LeBlanc, E.V. and C.C. Colpitts, The green tea catechin EGCG provides proof-of-concept for a pan-coronavirus attachment inhibitor. Sci Rep, 2022. 12(1): p. 12899.

47. Utani, A., et al., A unique sequence of the laminin alpha 3 G domain binds to heparin and promotes cell adhesion through syndecan-2 and -4. J Biol Chem, 2001. 276(31): p. 28779–88.

48. Wang, C., et al., Antigenic structure of the human coronavirus OC43 spike reveals exposed and occluded neutralizing epitopes. Nat Commun, 2022. 13(1): p. 2921.

49. Lind, T., et al., The putative tumor suppressors EXT1 and EXT2 are glycosyltransferases required for the biosynthesis of heparan sulfate. J Biol Chem, 1998. 273(41): p. 26265–8.

50. Brass, A.L., et al., The IFITM proteins mediate cellular resistance to influenza A H1N1 virus, West Nile virus, and dengue virus. Cell, 2009. 139(7): p. 1243–54.

51. Compton, A.A., et al., IFITM proteins incorporated into HIV-1 virions impair viral fusion and spread. Cell Host Microbe, 2014. 16(6): p. 736–47.

52. Zhao, X., et al., Interferon induction of IFITM proteins promotes infection by human coronavirus OC43. Proc Natl Acad Sci U S A, 2014. 111(18): p. 6756–61.

53. Jia, R., et al., The N-terminal region of IFITM3 modulates its antiviral activity by regulating IFITM3 cellular localization. J Virol, 2012. 86(24): p. 13697–707.

54. Bonifacino, J.S. and L.M. Traub, Signals for sorting of transmembrane proteins to endosomes and lysosomes. Annu Rev Biochem, 2003. 72: p. 395–447.

55. Tartell, M.A., et al., Interferon-gamma induced RNF213 targets nascent double membrane vesicles to inhibit coronavirus primary translation. bioRxiv, 2026: p. 2026.06.02.729663.

56. Orizio, F., et al., Human sialic acid acetyl esterase: Towards a better understanding of a puzzling enzyme. Glycobiology, 2015. 25(9): p. 992–1006.

57. Rogers, G.N., et al., Influenza C virus uses 9-O-acetyl-N-acetylneuraminic acid as a high affinity receptor determinant for attachment to cells. J Biol Chem, 1986. 261(13): p. 5947–51.

58. Mani, K., et al., The heparan sulfate-specific epitope 10E4 is NO-sensitive and partly inaccessible in glypican-1. Glycobiology, 2004. 14(7): p. 599–607.

59. Hassan, Z., et al., OC43 clinical isolate spike proteins have distinct carbohydrate-binding properties. Nat Commun, 2026. 17(1).

60. Busse, M., et al., Contribution of EXT1, EXT2, and EXTL3 to heparan sulfate chain elongation. J Biol Chem, 2007. 282(45): p. 32802–10.

61. Klein, S., et al., IFITM3 blocks influenza virus entry by sorting lipids and stabilizing hemifusion. Cell Host Microbe, 2023. 31(4): p. 616–633.e20.

62. Liu, G. and M.U. Gack, An Optimized Circular Polymerase Extension Reaction-based Method for Functional Analysis of SARS-CoV-2. bioRxiv, 2022.

